# Early candidate biomarkers in urine of Walker-256 lung metastasis rat model

**DOI:** 10.1101/306050

**Authors:** Jing Wei, Na Ni, Linpei Zhang, Youhe Gao

## Abstract

Cancer metastasis accounts for the majority of deaths by cancer. Detection of cancer metastasis at its early stage is important for the management and prediction of cancer progression. Urine, which is not regulated by homeostatic mechanisms, reflects systemic changes in the whole body and can potentially be used for the early detection of cancer metastasis. In this study, a lung metastasis of a Walker-256 rat model was established by tail-vein injection of Walker-256 cells. Urine samples were collected at days 2, 4, 6 and 9 after injection, and the urinary proteomes were profiled using liquid chromatography coupled with tandem mass spectrometry (LC-MS/MS). The urinary protein patterns changed significantly with the development of Walker-256 lung metastasis. On the fourth day, lung metastasis nodules appeared. On the sixth day, clinical symptoms started. On days 2, 4, 6 and 9, 11, 25, 34 and 44 differential proteins were identified in 7 lung metastatic rats by LC-MS/MS. Seventeen of these 62 differential proteins were identified on the second day, and 18 of them were identified on the fourth day. The differential urinary proteins changed significantly two days before lung metastasis nodules appeared. Differential urinary proteins differed in Walker-256 lung metastasis rat models and Walker-256 subcutaneous rat models. A total of 9 differential proteins (NHRF1, CLIC1, EZRI, AMPN, ACY1A, HSP7C, BTD, NID2, and CFAD) were identified in 7 lung metastatic rats at one or more common time points, and these 9 differential proteins were not identified in the subcutaneous rat model. Seven of these 9 differential proteins were associated with both breast cancer and lung cancer, eight of the nine were identified on the second day, and 8 of the nine can be identified on the fourth day; these early changes in urine were also identified with differential abundances at late stages of lung metastasis. Our results indicate that (1) the urine proteome changed significantly, even on the second day after tail-vein injection of Walker-256 cells and that (2) the urinary differential proteins were different in Walker-256 lung metastatic tumors and Walker-256 subcutaneous tumors. Our results provide the potential to detect early breast cancer lung metastasis, monitor its progression and differentiate it from the same cancer cells grown at other locations.

## Introduction

Cancer metastasis is a process in which cancer cells are disseminated from primary tumor tissue to different sites through blood vessels and lymphatic vessels. Lung, brain, bone and liver are the common metastatic organs in cancer patients[1]. Distant organ metastasis accounts for most cancer morbidity and mortality and nearly 90% of cancer death[2] and is usually accompanied by a poor 5-year survival rate as well as limited treatment strategies[3]. Due to special lung-specific immunoregulatory mechanisms, tumor colonization occurs more readily in an immunologically permissive environment[4]. Therefore, many cancer metastases such those of as breast cancer and malignant melanoma occur more easily in lung. The early detection of cancer metastasis is still elusive, as finding and predicting specific distant metastatic organs is difficult, especially in early-stage cancer without clinical symptoms. Therefore, the early detection of cancer metastasis can significantly improve the survival rate and effective therapies of cancer patients and also helps in monitoring cancer metastasis progression in time.

Biomarkers are measurable changes associated with the physiological or pathophysiological processes of disease and usually derive from tissue, blood and tumor cells [5]. Because of the homeostatic mechanisms in the internal environment, the levels of important factors in blood tend to be stable to protect the stability of the internal environment[6]. In addition, without the control of homeostatic mechanisms, urine can accumulate systemic changes from the whole body and thus has the potential to reflect the small pathophysiological changes of disease[7]. Therefore, urine has the potential to reflect early changes in disease. However, whether time-course analyses of urine proteins can reveal reliable cancer metastasis biomarkers at different stages of cancer metastasis is unclear, as urinary proteins are easily affected by complicated factors such as medicine and diet, especially in human samples. Therefore, using a small number of animal models can help to determine the direct relationship between urine protein changes and related diseases such as cancer metastasis because the genetic and environmental factors are minimized [8]. In addition, determining an exact description of cancer metastasis is very helpful for the identification of cancer metastasis biomarkers, especially in early stages.

Various studies have applied urinary proteomics to discover cancer biomarkers for the early diagnosis and monitoring of cancer[9–11]. However, most of these studies used clinical urine samples from breast cancer patients who had already had distant metastases to viscera or bone[10]. It is difficult to clinically collect the exact early stages of breast cancer lung metastasis samples. Using animal models renders the exact starting point of cancer lung metastasis available, which is very helpful in the identification of biomarkers in the early stage of cancer lung metastasis.

Walker-256 cells are mammary gland carcinoma cells[12], and the Walker-256 lung carcinoma metastasis rat model is a well-known cancer lung metastasis rat model for studies of lung metastasis progression, such as evaluating the effects of some drugs on the development of Walker-256 lung metastases [13]. In this study, the Walker-256 lung carcinoma metastasis rat model was established by tail-vein injection of Walker-256 tumor cells. Urine samples were collected from lung metastasis rat models on days 2, 4, 6, and 9 for further urine proteome analysis. By comparing the differential proteins of Walker-256 lung carcinoma metastasis rats and subcutaneous rats[9], early lung metastasis associated urine biomarkers was identified. The workflow of this research is presented in Figure 1.

**Figure 1.**
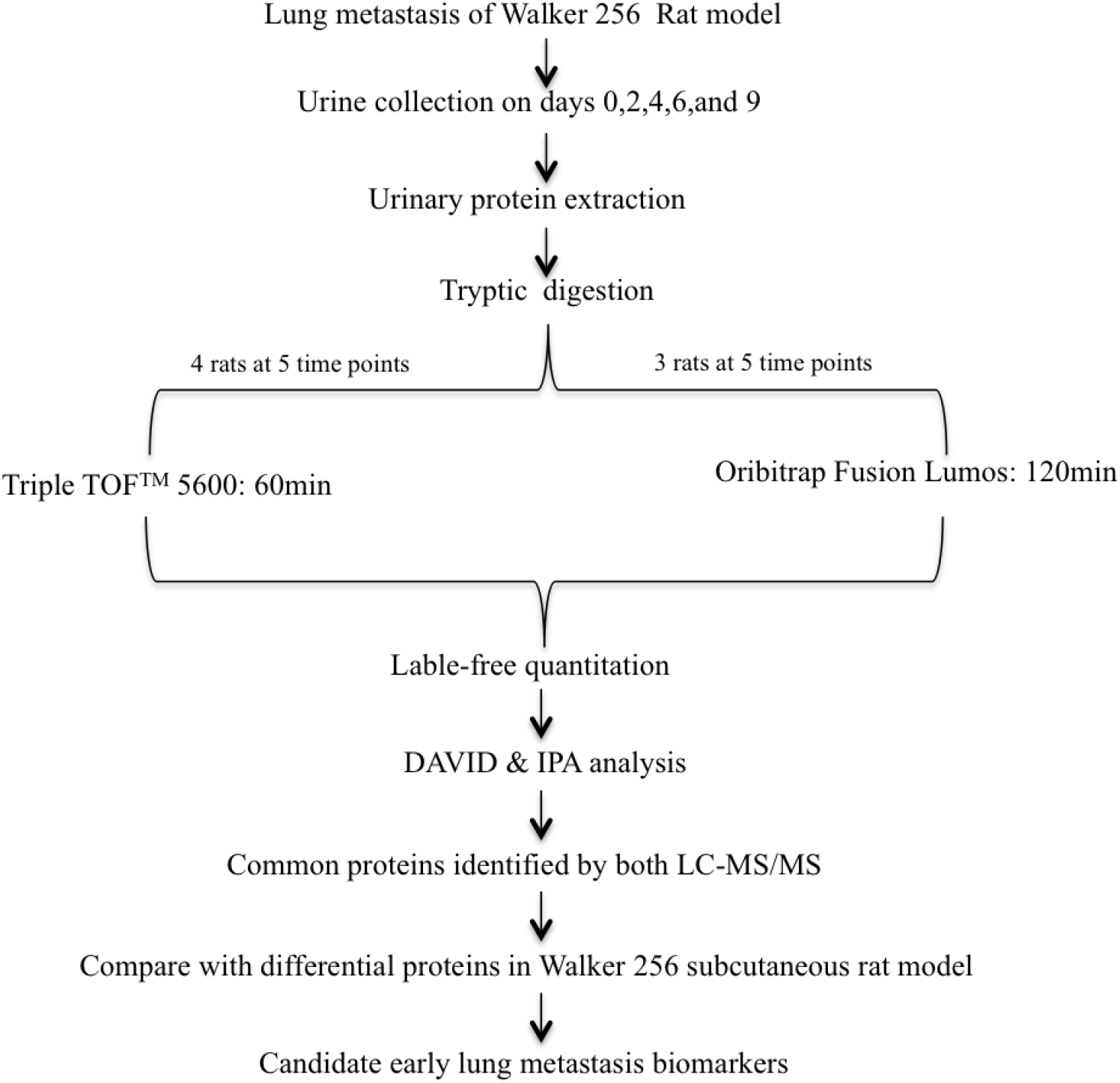
Workflow of urinary proteomic discovery in this study. Urine samples were collected on days 2, 4, 6, and 9.

## Materials and methods

### Experimental animals

Male Wistar rats (150 ±20 g) and Sprague-Dawley (SD) rats (70 ± 20 g) were purchased from the Beijing Vital River Laboratory Animal Technology Co, Ltd. All animals were housed with free access to a standard laboratory diet and water with controlled indoor temperature (22 ±1°C) and humidity (65 ~ 70%) and a 12 h/12 h light-dark cycle. All animal protocols governing the experiments in this study were approved by the Institute of Basic Medical Sciences Animal Ethics Committee, Peking Union Medical College (Approved ID: ACUC-A02-2014-008). The study was performed according to the guidelines developed by the Institutional Animal Care and Use Committee of Peking Union Medical College. All efforts were made to minimize suffering.

### Rat model establishment

The Walker-256 lung carcinoma metastasis rat model was established as reported previously[13]. The Walker-256 carcinosarcoma cells were purchased from the Cell Culture Center of the Chinese Academy of Medical Sciences (Beijing, China). Briefly, male SD rats were used for ascitic tumor cell cultivation. After two cell passages, the Walker-256 tumor cells were collected, centrifuged, and resuspended in phosphate-buffered saline (PBS) for the following establishment of rat models. The cell viability was assessed by the trypan blue exclusion test. Walker-256 cells were stained with 0.4% trypan blue solution and then counted using a hemocytometer. Their viability was approximately 95% before the tail-vein injection.

Male Wistar rats were randomly divided into two groups: the Walker-256 lung carcinoma metastasis group (n = 10) and the control group (n = 4). The Walker-256 lung carcinoma metastasis group was injected with 2×10^6^ viable Walker-256 cells in 100 *μ*L of PBS by tail-vein injection. The control group was tail-vein injected with the same volume of PBS. The animals were anesthetized with sodium pentobarbital solution (4 mg/kg) before the tail-vein injection.

### Lung histopathology

For histopathology, rats were sacrificed on days 2, 4, 6, and 9 by using an overdose of sodium pentobarbital anesthetic. The whole lung tissue was fixed in 4% formalin fixative and embedded in paraffin.

Then, the paraffin sections (4-*μ*m thick) were stained with hematoxylin and eosin (HE) to reveal the metastatic nodules.

### Urine collection and protein extraction

First, the rats were accommodated in metabolic cages for 2-3 days for urine sample colleting. Then, the urine samples were collected from each rat (from either the lung metastasis group or the control group) on days 2, 4, 6, and 9 after Walker-256 cell or PBS tail-vein injection. Each rat was placed in metabolic cages with free access to water and without food to avoid contamination overnight for the collection of the urine samples in the following 12 h.

Urine samples were centrifuged at 12,000 g for 30 min at 4°C immediately to remove cell debris. Then, the supernatants were precipitated with three volumes of ethanol at -20°C for 2 h. After centrifugation at 12,000 g for 30 min, the pellets were resuspended in lysis buffer (8 mol/L urea, 2 mol/L thiourea, 50 mmol/L Tris, and 25 mmol/L DTT) at 4°C for 2 h. Finally, after centrifugation at 4°C and 12, 000 g for 30 min, the supernatants of each sample were measured by using the Bradford assay. The protein samples were stored at −80°C for later use.

### SDS-PAGE analysis

After Walker-256 cell tail-vein injection, 25 μg of protein from each sample on days 2, 4, 6, and 9 and the control group on day 2 was added to loading buffer (50 mmol/L Tris-HCl, pH 6.8, 50 mol/L DTT, 0.5% SDS, and 10% glycerol). Then, all these protein samples were incubated at 98°C for 10 min. The urine protein samples from randomly selected Walker-256 lung carcinoma metastasis rats were resolved by 12% sodium dodecyl sulfate-polyacrylamide gel electrophoresis (SDS-PAGE).

### Protein digestion and peptide preparation

Urine protein samples of Walker-256 lung carcinoma metastasis rats on days 2, 4, 6, and 9 and the control group on day 2 were randomly selected for proteomic analysis. Proteins were digested with trypsin (Trypsin Gold, Mass Spec Grade, Promega, Fitchburg, Wisconsin, USA) by using filter-aided sample preparation methods, as reported previously[14]. Briefly, 100 *μ*g of proteins were loaded onto 10-kD cutoff filter devices (Pall, Port Washington, NY) and washed with UA (8 M urea in 0.1 M Tris-HCl, pH 8.5) at 14,000 g for 40 min at 18°C twice. Then, 25 mmol/L NH_4_HCO_3_ was added to wash the protein. Each urinary protein was subsequently denatured with 20 mM DTT at 37°C for 1 h and then alkylated with 50 mM iodoacetamide (IAA) for 40 min in the dark. After being washed twice with UA and 3 times with 25 mmol/L NH_4_HCO_3_, the denatured proteins were resuspended with 25 mmol/L NH_4_HCO_3_ and digested with trypsin (enzyme to protein ratio of 1:50) at 37°C for 12-16 h. Finally, the collected peptide mixtures were desalted using Oasis HLB cartridges (Waters, Milford, MA) and then dried by vacuum evaporation (Thermo Fisher Scientific, Bremen, Germany).

### LC-MS/MS analysis

Digested peptides were re-dissolved in 0.1% formic acid to a concentration of 0.5 *μ*g/*μ*L. Then, 2 *μ*g of each sample was transferred to a reversed-phase microcapillary column using a Waters ultra-performance liquid chromatography (UPLC) system, and peptides were separated on a 10-cm fused silica column. The elution from the fused silica column was performed in 60 min with a gradient of 5%–28% buffer B (0.1% formic acid and 99.9% acetonitrile (ACN); flow rate, 0.3 *μ*L/min). The peptides were analyzed using an AB SCIEX (Framingham, MA, US) Triple TOF 5600 mass spectrometry (MS) system. Samples from four Walker-256 lung carcinoma metastasis rats at 4 time points and four control rats were randomly chosen for this study. Each sample was analyzed twice. Digested peptides were re-dissolved in 0.1% formic acid to a concentration of 0.5 *μ*g/*μ*L. For analysis, 1 *μ*g of each peptide from an individual sample was loaded onto a trap column and separated on a reverse-phase C18 column (50 *μ*m × 150 mm, 2 *μ*m) using the EASY-nLC 1200 HPLC system (Thermo Fisher Scientific, Waltham, MA). The elution for the analytical column lasted 120 min at a flow rate of 300 nL/min. Then, the peptides were analyzed with an Orbitrap Fusion Lumos Tribrid mass spectrometer (Thermo Fisher Scientific, Waltham, MA). MS data were acquired in high-sensitivity mode using the following parameters: data-dependent MS/MS scans per full scan with top-speed mode (3 s), MS scans at a resolution of 120,000 and MS/MS scans at a resolution of 30,000 in the Orbitrap, 30% HCD collision energy, charge-state screening (+2 to +7), dynamic exclusion (exclusion duration 30 s), and a maximum injection time of 45 ms.

### Database searching and label-free quantitation

The MS/MS data of Walker-256 lung metastasis rat samples were searched using Mascot software (version 2.4.1, Matrix Science, London, UK) against the SwissProt rat database (released in February 2017, containing 7992 sequences). For Triple TOF 5600^TM^, the parameters were set as follows: the fragment ion mass tolerance was set to 0.05 Da, and the parent ion tolerance was set to 0.05 Da. The search allowed two missed cleavage sites in the trypsin digestion. The carbamidomethylation of cysteines was considered a fixed modification, and the oxidation of methionine and deamidation of asparagine were considered variable modifications. For the Orbitrap Fusion Lumos, the parent ion tolerance was set to 10 ppm, and the fragment ion mass tolerance was set to 0.02 Da. Carbamidomethylation of cysteine was set as a fixed modification, and the oxidation of methionine was considered a variable modification. The specificity of trypsin digestion was set for cleavage after K or R, and two missed trypsin cleavage sites were allowed.

Peptide and protein identification was further validated by Progenesis LC-MS/MS software (version 4.1, Nonlinear, Newcastle upon Tyne, UK) and Scaffold (version 4.7.5, Proteome Software Inc., Portland, OR). For Progenesis, the acquired data from the MS scans were transformed and stored in peak lists using a proprietary algorithm. Features with only one charge or more than five charges were excluded from the analyses. Protein abundance was calculated from the sum of all unique peptide ion abundances for a specific protein in each run. The normalization of abundances was required to allow comparisons across different sample runs by this software. For further quantitation, all peptides of an identified protein were included. Proteins identified by more than one peptide were retained. For Scaffold, peptide identifications were accepted at a false discovery rate (FDR) of less than 1.0% by the Scaffold Local FDR algorithm, and protein identifications were accepted at an FDR less than 1.0% with at least two unique peptides. Comparisons across different samples were performed after normalization of total spectra using Scaffold software. Spectral counting was used to compare protein abundances at different time points according to a previously described procedure.

### Gene ontology and ingenuity pathway analysis

All proteins identified to be differentially expressed between the control and lung carcinoma metastasis rats were assigned a gene symbol using DAVID [15] and analyzed by Gene Ontology (GO) based on biological process, cellular component and molecular function categories. The biological pathway analysis of differential proteins analyzed at four time points were performed by IPA software (Ingenuity Systems, Mountain View, CA, USA)

### Statistical analysis

Statistical analysis was performed with GraphPad Prism version 7.0 (GraphPad, San Diego, CA). Comparisons between data from samples of four time points were conducted using repeated-measures one-way ANOVA followed by multiple comparison analysis with the least significant difference (LSD) test. Group differences resulting in *P* < 0.05 were considered statistically significant.

## Results and discussion

### Body weight changes of Walker-256 lung carcinoma metastasis rat model

From 6 days after the tail-vein injection of Walker-256 cells, the average body weight of lung carcinoma metastasis rats was lower than that of the control rats (Figure 2), and reduced food intake was also observed in lung carcinoma metastasis rats. On day 9 after the tail-vein injection of Walker-256 cells, the body weight of lung carcinoma metastasis rats was significantly lower than that of the control group. Therefore, we believed that days 2 and 4 were early time points during Walker-256 lung metastasis.

**Figure 2.**
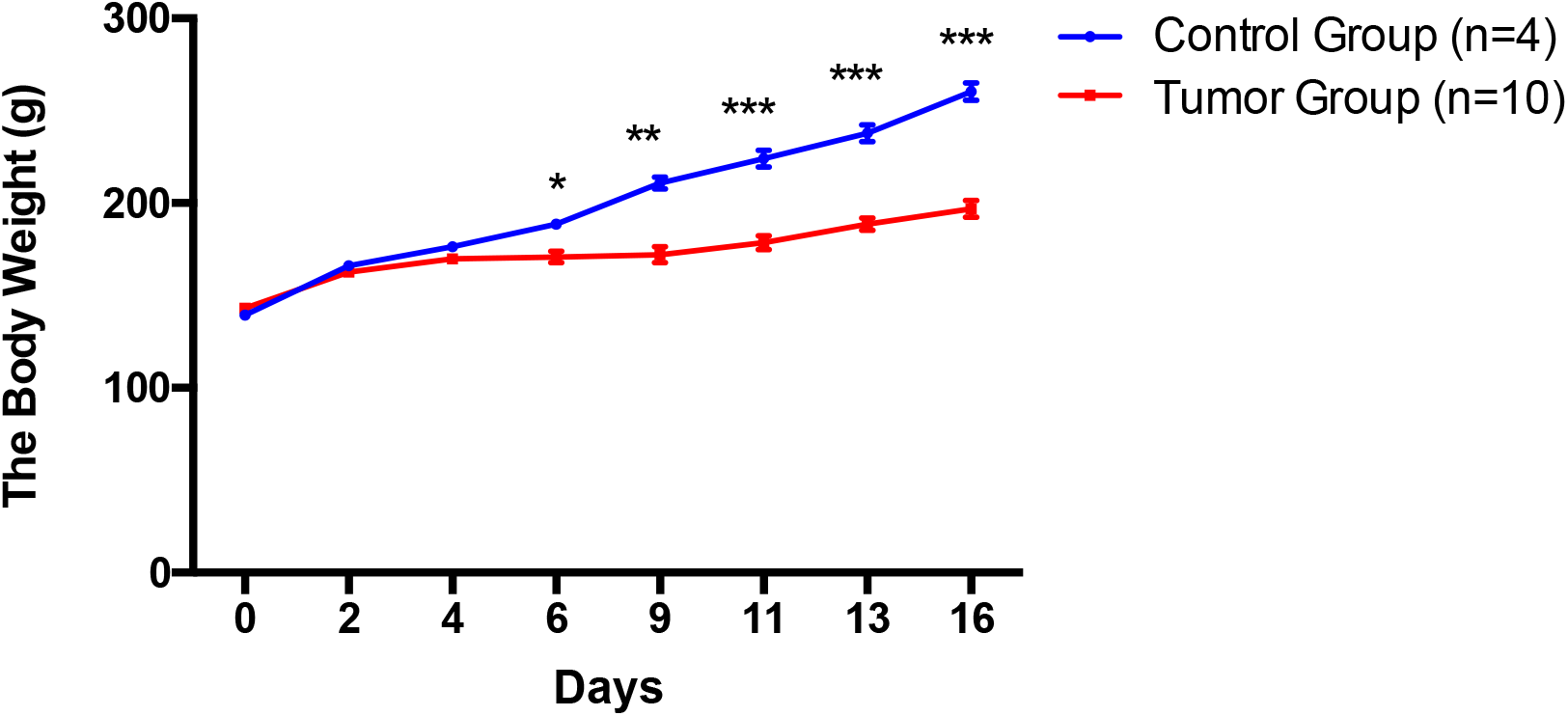
The body weight changes of Walker-256 lung carcinoma metastasis rats.

### Pathological changes in Walker-256 lung metastasis rat model

The pathological changes in the Walker-256 lung carcinoma metastasis rats at different time points are shown in Figure 3. The lung metastasis nodules appeared on day 4, and their number and volumes increased in Walker-256 lung carcinoma metastasis rats during lung metastasis progression. In addition, the metastatic Walker-256 cells arranged closely in lung metastatic rats, and the majority of cells showed round or elliptic morphologies accompanied by poor differentiation, while their nuclei were large, irregular, and hyperchromatic. More importantly, the lung metastasis nodules were scattered throughout the lung parenchyma, while the invasion of Walker-256 cells destroyed the lung tissue structure and the alveolar structure.

**Figure 3.**
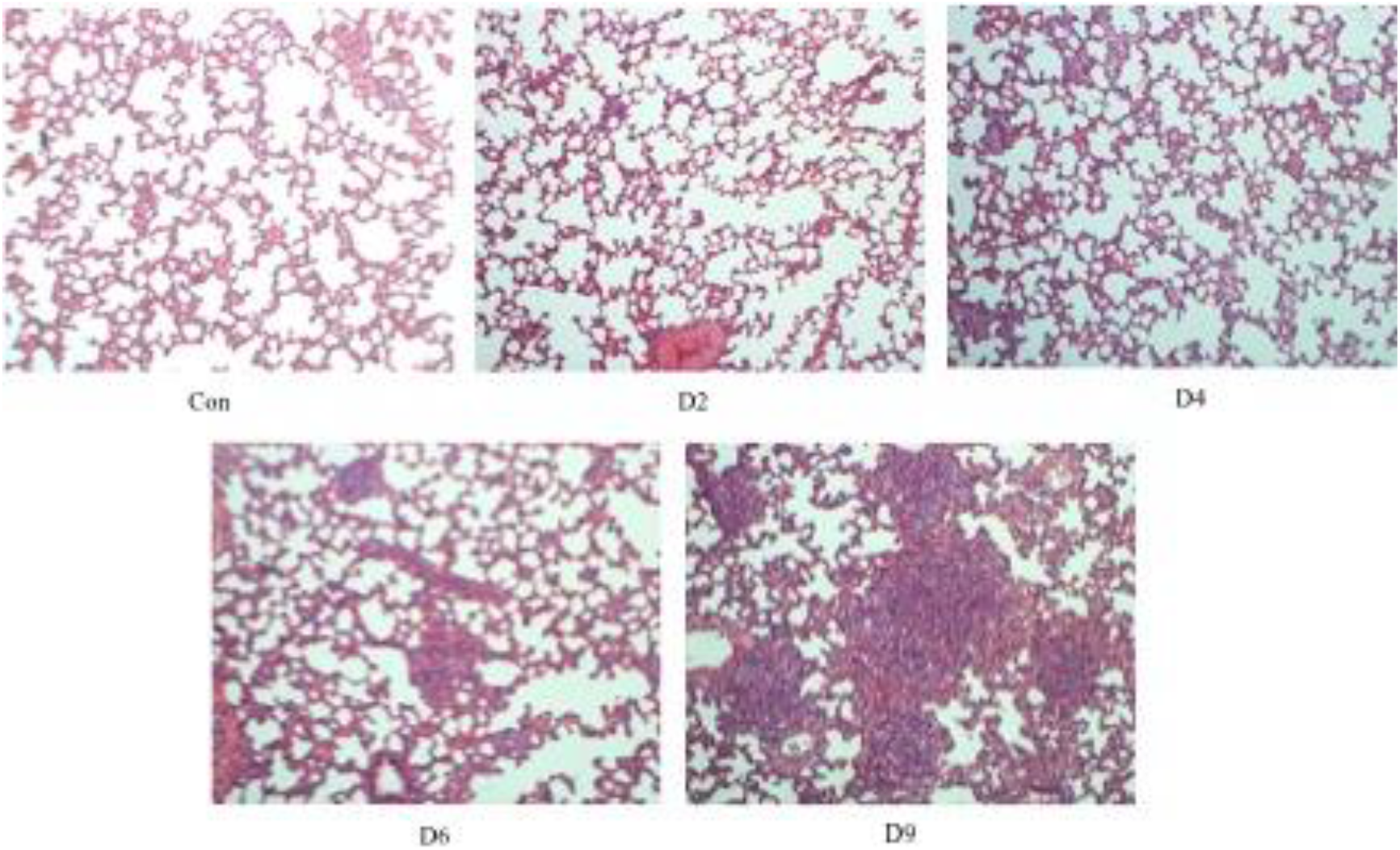
Pathological changes in Walker-256 lung metastatic rats. The magnification was 100× for the images of H&E staining.

### SDS-PAGE of Walker-256 lung metastasis rat model

Urine samples collected at different time points from Walker-256 lung carcinoma metastasis rats were separated by 12% SDS-PAGE. As shown in Figure 4, the patterns in urine sample proteins from a representative Walker-256 lung metastasis rat changed significantly during lung metastatic progression (days 1, 2, 4, 6, 9, 11, 13 and 16). Similar patterns were observed in another 6 rats, suggesting consistent lung metastatic progression in the chosen rats. It can be seen in Figure 4 that on days 6 and 9, the protein band intensities increased significantly, which is consistent with the times at which the body weight changed.

**Figure 4.**
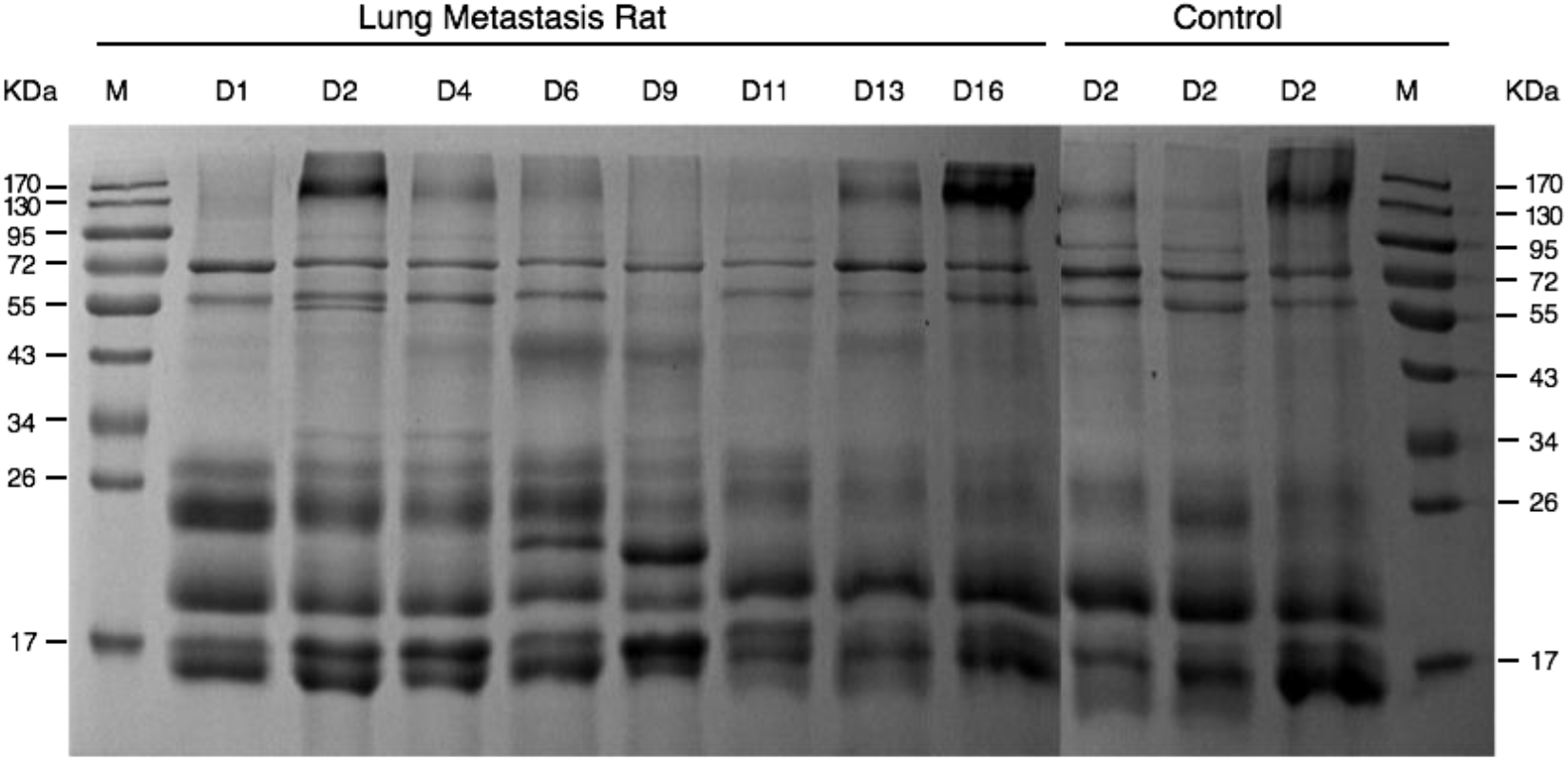
Dynamic protein patterns in the urine of Walker-256 lung carcinoma metastasis rats.

### The urine proteome was significantly different in the Walker-256 lung metastasis rat model

A total of 263 urinary proteins were identified with at least two peptides by using a Triple TOF 5600^TM^ mass spectrometer, and 839 urinary proteins were identified with <1% FDR at the protein level with at least two peptides by using Orbitrap Fusion Lumos. The differential proteins were screened by the following criteria: fold change ≥1.5 or ≤ 0.67, confidence score ≥ 200, *P* < 0.05 for differences in the protein level between the metastasis rat model and the control group, protein spectral counts or the normalized abundance from every rat in the high-abundance group greater than those in the low-abundance group, and the average spectral count in the high-abundance group ≥ 4. By using these screening criteria, 102 differential proteins were identified by using the Triple TOF 5600^TM^ mass spectrometer, and 171 differential proteins were identified by using the Orbitrap Fusion Lumos.

The overlap of differential proteins identified at different stages in 7 lung metastatic rats is shown by a Venn diagram (Figure 5). There were 85, 81, 142, and 133 differential proteins on days 2, 4, 6, and 9 after tail-vein injection of Walker-256 cells, respectively. In addition, 11, 25, 34, and 44 proteins that were differentially expressed in all 7 lung metastatic rats were identified by two mass spectrometers on days 2, 4, 6, and 9, respectively. There were 62 differential proteins that differed at one or more of the same time points and were identified by using two mass spectrometers. Among these 62 differential proteins, the levels of 29 differential urinary proteins changed significantly on the second day and before lung metastasis nodules appeared, indicating their potential roles in the early detection of lung cancer metastasis (Table 1). A data processing flowchart is presented in Figure 6.

**Figure 5.**
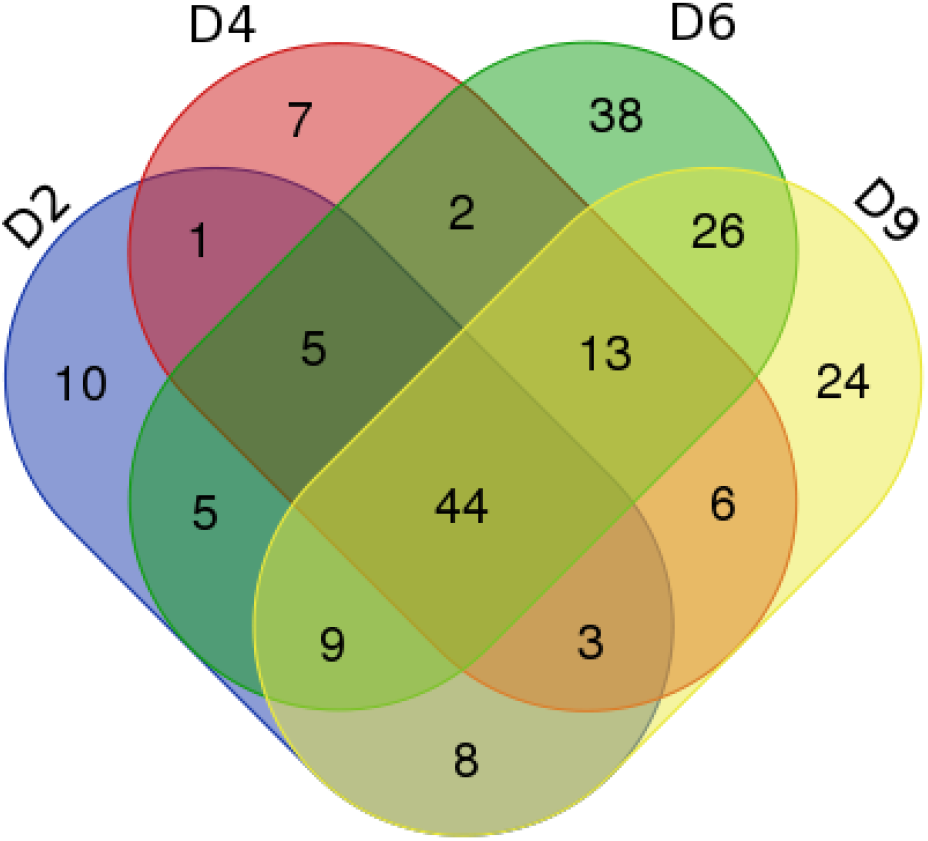
Evaluation of the overlap of differential proteins identified at different metastatic phases in 7 lung metastatic rats.

**Figure 6.**
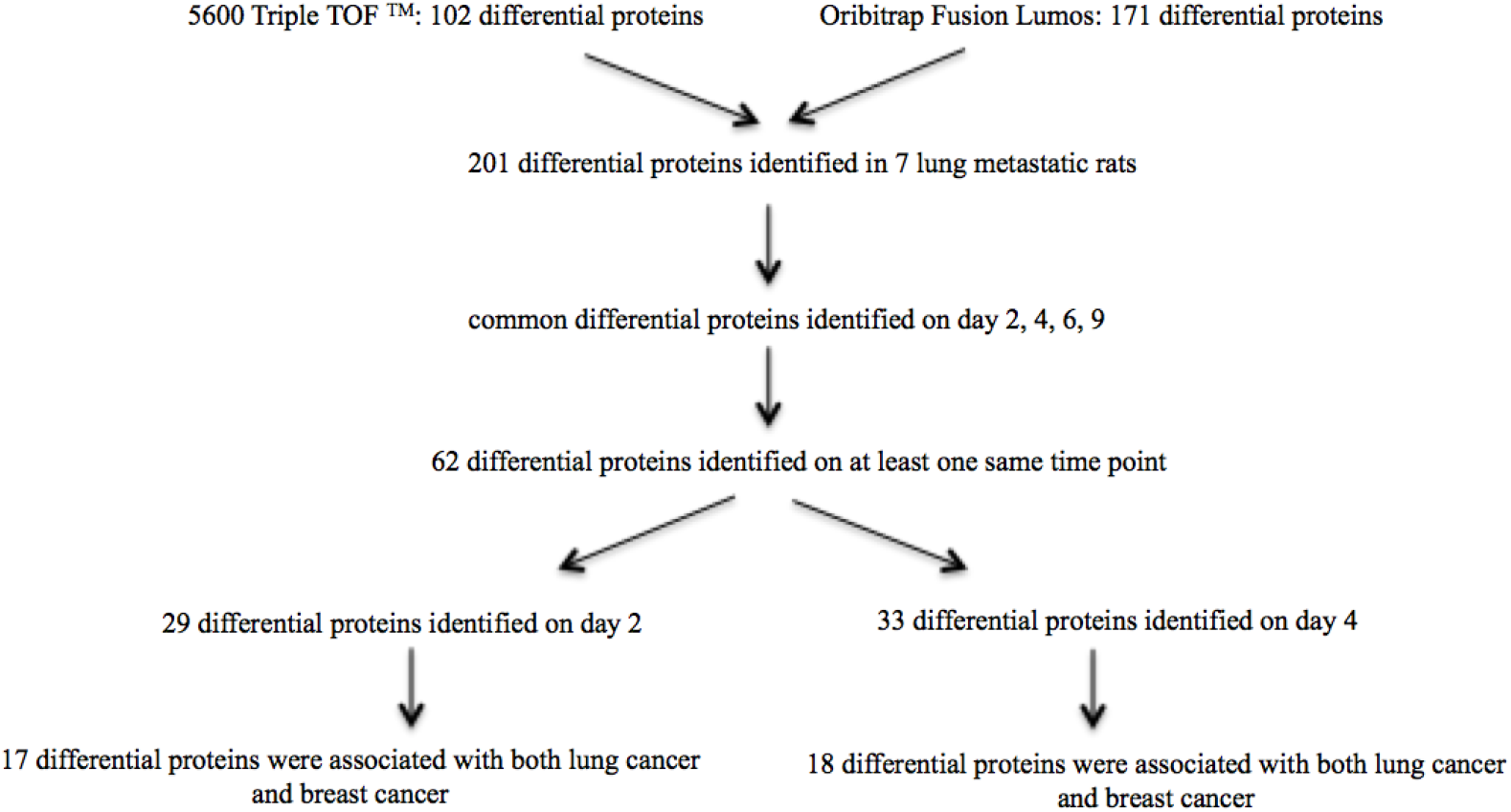
Steps for processing the data from 7 lung metastatic rats.

Some of these 29 differential proteins have been reported to be clinical lung cancer biomarkers and also associated with breast cancer metastasis. For example, (1) galectin-3-binding protein (LG3BP) is a candidate biomarker for the diagnosis of large-cell neuroendocrine lung carcinoma[16], and it can also induce galectin-mediated tumor cell aggregation to increase the survival of cancer cells in the bloodstream during the metastasis [17]. As another example, (2) neutrophil gelatinase-associated lipocalin (NGAL) is a potential biomarker for early stages of lung tumorigenesis[18]. Additionally, the stroma-secreted NGAL promotes breast cancer metastasis in vitro and in vivo, thereby contributing to tumor progression[19]. (3) The baseline soluble intercellular adhesion molecule 1 (ICAM-1) levels in serum were evaluated as an additional prognostic factor in patients with small cell lung cancer (SCLC). In addition, the soluble protein ICAM-1 may also be a predictive marker for an objective response during chemotherapy for patients with extensive disease (p = 0.001)[20]. In breast cancer, ICAM-1 can also activate intracellular signaling pathways in cancer cells, leading to enhanced cell motility, invasion and metastasis[21]. (4) In advanced non-small cell lung cancer (NSCLC) patients, the pretreatment serum C-reactive protein (CRP) was associated with a poor outcome of treatment with pemetrexed[22]. In addition, a positive association between pre-diagnostic high-sensitivity CRP (hs-CRP) was reported with breast cancer risk[23]. (5) Apolipoprotein E (ApoE) levels significantly increased in the pleural effusion of patients with NSCLC, which serves as a potential marker for the diagnosis of malignant pleural effusions (MPEs) as well as the differential diagnosis of MPE in NSCLC[24]. Additionally, apolipoprotein E expression promoted lung adenocarcinoma proliferation and migration, which can be a potential survival marker in lung cancer[25]. In breast cancer, the serum level of apolipoprotein E was also reported to correlate with disease progression and poor prognosis[26]. Other differential proteins are annotated in Table 1.

According to our results, we suggest that it is more appropriate to use a protein panel for a biomarker, as the specificities of single protein biomarkers are not significant enough.

**Table 1.**
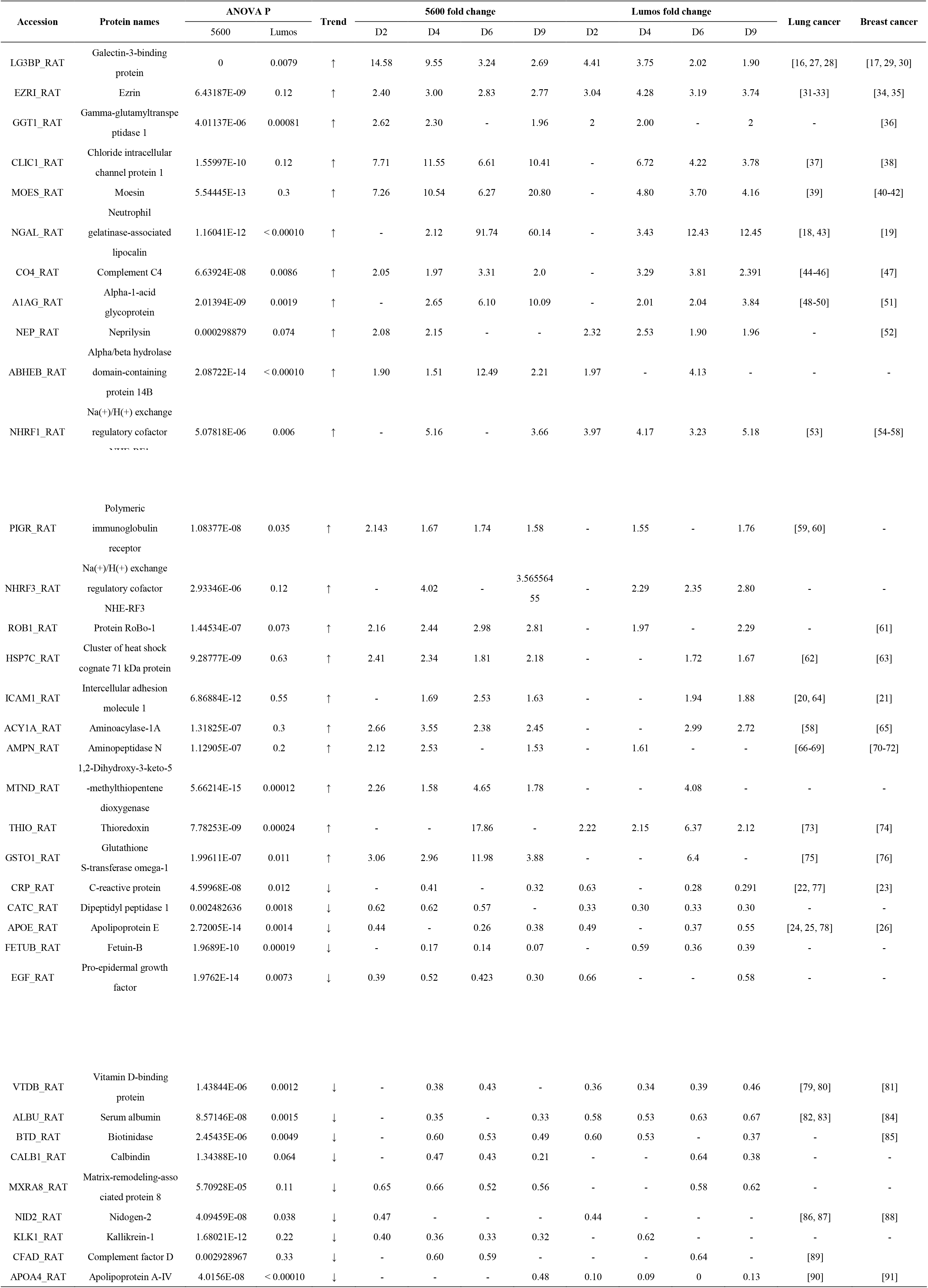
The differential proteins identified at days 2 and 4 in 7 lung metastatic rats by using two mass spectrometers.

| Accession | Protein names | ANOVA P |  | Trend | 5600 fold change |  |  |  | Lumos fold change |  |  |  | Lung cancer | Breast cancer |
| --- | --- | --- | --- | --- | --- | --- | --- | --- | --- | --- | --- | --- | --- | --- |
|  |  | 5600 | Lumos |  | D2 | D4 | D6 | D9 | D2 | D4 | D6 | D9 |  |  |
| LG3BP_RAT | Galectin-3-binding protein | 0 | 0.0079 | ↑ | 14.58 | 9.55 | 3.24 | 2.69 | 4.41 | 3.75 | 2.02 | 1.90 | [16, 27, 28] | [17, 29, 30] |
| EZRI_RAT | Ezrin | 6.43187E-09 | 0.12 | ↑ | 2.40 | 3.00 | 2.83 | 2.77 | 3.04 | 4.28 | 3.19 | 3.74 | [31-33] | [34, 35] |
| GGT1_RAT | Gamma-glutamyltranspeptidase 1 | 4.01137E-06 | 0.00081 | ↑ | 2.62 | 2.30 | - | 1.96 | 2 | 2.00 | - | 2 | - | [36] |
| CLIC1_RAT | Chloride intracellular channel protein 1 | 1.55997E-10 | 0.12 | ↑ | 7.71 | 11.55 | 6.61 | 10.41 | - | 6.72 | 4.22 | 3.78 | [37] | [38] |
| MOES_RAT | Moesin | 5.54445E-13 | 0.3 | ↑ | 7.26 | 10.54 | 6.27 | 20.80 | - | 4.80 | 3.70 | 4.16 | [39] | [40-42] |
| NGAL_RAT | Neutrophil gelatinase-associated lipocalin | 1.16041E-12 | < 0.00010 | ↑ | - | 2.12 | 91.74 | 60.14 | - | 3.43 | 12.43 | 12.45 | [18, 43] | [19] |
| CO4_RAT | Complement C4 | 6.63924E-08 | 0.0086 | ↑ | 2.05 | 1.97 | 3.31 | 2.0 | - | 3.29 | 3.81 | 2.391 | [44-46] | [47] |
| A1AG_RAT | Alpha-1-acid glycoprotein | 2.01394E-09 | 0.0019 | ↑ | - | 2.65 | 6.10 | 10.09 | - | 2.01 | 2.04 | 3.84 | [48-50] | [51] |
| NEP_RAT | Neprilysin | 0.000298879 | 0.074 | ↑ | 2.08 | 2.15 | - | - | 2.32 | 2.53 | 1.90 | 1.96 | - | [52] |
| ABHEB_RAT | Alpha/beta hydrolase domain-containing protein 14B | 2.08722E-14 | < 0.00010 | ↑ | 1.90 | 1.51 | 12.49 | 2.21 | 1.97 | - | 4.13 | - | - | - |
| NHRF1_RAT | Na(+)/H(+) exchange regulatory cofactor NHE-RF1 | 5.07818E-06 | 0.006 | ↑ | - | 5.16 | - | 3.66 | 3.97 | 4.17 | 3.23 | 5.18 | [53] | [54-58] |
| PIGR_RAT | Polymeric<br>immunoglobulin<br>receptor | 1.08377E-08 | 0.035 | ↑ | 2.143 | 1.67 | 1.74 | 1.58 | - | 1.55 | - | 1.76 | [59, 60] | - |
| NHRF3_RAT | Na(+)/H(+) exchange<br>regulatory cofactor<br>NHE-RF3 | 2.93346E-06 | 0.12 | ↑ | - | 4.02 | - | 3.565564<br>55 | - | 2.29 | 2.35 | 2.80 | - | - |
| ROB1_RAT | Protein RoBo-1 | 1.44534E-07 | 0.073 | ↑ | 2.16 | 2.44 | 2.98 | 2.81 | - | 1.97 | - | 2.29 | - | [61] |
| HSP7C_RAT | Cluster of heat shock<br>cognate 71 kDa protein | 9.28777E-09 | 0.63 | ↑ | 2.41 | 2.34 | 1.81 | 2.18 | - | - | 1.72 | 1.67 | [62] | [63] |
| ICAM1_RAT | Intercellular adhesion<br>molecule 1 | 6.86884E-12 | 0.55 | ↑ | - | 1.69 | 2.53 | 1.63 | - | - | 1.94 | 1.88 | [20, 64] | [21] |
| ACY1A_RAT | Aminoacylase-1A | 1.31825E-07 | 0.3 | ↑ | 2.66 | 3.55 | 2.38 | 2.45 | - | - | 2.99 | 2.72 | [58] | [65] |
| AMPN_RAT | Aminopeptidase N | 1.12905E-07 | 0.2 | ↑ | 2.12 | 2.53 | - | 1.53 | - | 1.61 | - | - | [66-69] | [70-72] |
| MTND_RAT | 1,2-Dihydroxy-3-keto-5-<br>-methylthiopentene<br>dioxygenase | 5.66214E-15 | 0.00012 | ↑ | 2.26 | 1.58 | 4.65 | 1.78 | - | - | 4.08 | - | - | - |
| THIO_RAT | Thioredoxin | 7.78253E-09 | 0.00024 | ↑ | - | - | 17.86 | - | 2.22 | 2.15 | 6.37 | 2.12 | [73] | [74] |
| GSTO1_RAT | Glutathione<br>S-transferase omega-1 | 1.99611E-07 | 0.011 | ↑ | 3.06 | 2.96 | 11.98 | 3.88 | - | - | 6.4 | - | [75] | [76] |
| CRP_RAT | C-reactive protein | 4.59968E-08 | 0.012 | ↓ | - | 0.41 | - | 0.32 | 0.63 | - | 0.28 | 0.291 | [22, 77] | [23] |
| CATC_RAT | Dipeptidyl peptidase 1 | 0.002482636 | 0.0018 | ↓ | 0.62 | 0.62 | 0.57 | - | 0.33 | 0.30 | 0.33 | 0.30 | - | - |
| APOE_RAT | Apolipoprotein E | 2.72005E-14 | 0.0014 | ↓ | 0.44 | - | 0.26 | 0.38 | 0.49 | - | 0.37 | 0.55 | [24, 25, 78] | [26] |
| FETUB_RAT | Fetuin-B | 1.9689E-10 | 0.00019 | ↓ | - | 0.17 | 0.14 | 0.07 | - | 0.59 | 0.36 | 0.39 | - | - |
| EGF_RAT | Pro-epidermal growth<br>factor | 1.9762E-14 | 0.0073 | ↓ | 0.39 | 0.52 | 0.423 | 0.30 | 0.66 | - | - | 0.58 | - | - |
| VTDB_RAT | Vitamin D-binding protein | 1.43844E-06 | 0.0012 | ↓ | - | 0.38 | 0.43 | - | 0.36 | 0.34 | 0.39 | 0.46 | [79, 80] | [81] |
| ALBU_RAT | Serum albumin | 8.57146E-08 | 0.0015 | ↓ | - | 0.35 | - | 0.33 | 0.58 | 0.53 | 0.63 | 0.67 | [82, 83] | [84] |
| BTD_RAT | Biotinidase | 2.45435E-06 | 0.0049 | ↓ | - | 0.60 | 0.53 | 0.49 | 0.60 | 0.53 | - | 0.37 | - | [85] |
| CALB1_RAT | Calbindin | 1.34388E-10 | 0.064 | ↓ | - | 0.47 | 0.43 | 0.21 | - | - | 0.64 | 0.38 | - | - |
| MXRA8_RAT | Matrix-remodeling-associated protein 8 | 5.70928E-05 | 0.11 | ↓ | 0.65 | 0.66 | 0.52 | 0.56 | - | - | 0.58 | 0.62 | - | - |
| NID2_RAT | Nidogen-2 | 4.09459E-08 | 0.038 | ↓ | 0.47 | - | - | - | 0.44 | - | - | - | [86, 87] | [88] |
| KLK1_RAT | Kallikrein-1 | 1.68021E-12 | 0.22 | ↓ | 0.40 | 0.36 | 0.33 | 0.32 | - | 0.62 | - | - | - | - |
| CFAD_RAT | Complement factor D | 0.002928967 | 0.33 | ↓ | - | 0.60 | 0.59 | - | - | - | 0.64 | - | [89] |  |
| APOA4_RAT | Apolipoprotein A-IV | 4.0156E-08 | < 0.00010 | ↓ | - | - | - | 0.48 | 0.10 | 0.09 | 0 | 0.13 | [90] | [91] |

### Functional analysis of differential urine proteins in Walker-256 lung carcinoma metastasis rats

The functional annotation of differential proteins was performed by using DAVID[15]. Differential proteins at different metastatic time points were classified into biological process, molecular function, and molecular components (Figure 7). In the biological processes, epithelial cell differentiation, the regulation of immune system processes, and classical complement activation pathway were overrepresented on days 2, 4, 6 and 9. The ERK1 and ERK2 cascade was overrepresented on days 2, 4 and 6. The innate immune response and transport were overrepresented on days 4, 6 and 9. The cell adhesion was overrepresented on days 6 and 9. Interestingly, proteins representing the B cell receptor signaling pathway, the defense response to bacteria and the positive regulation of B cell activation appeared on day 9 (Figure 7A). The majority of these biological processes were reported to be associated with breast cancer metastasis or lung cancer. For example, the increasing levels of ERK1 and ERK2 were associated with breast cancer initiation, growth, and metastasis[92]. The persistent complement activation was reported for tumor cells in breast cancer, which consistent with the timing of its overrepresentation in this study[93]. The transport and cell adhesion processes were both overrepresented on days 6 and 9, which indicated the severe metastasis during lung tumor progression. Interestingly, on day 9, proteins representing the B cell activation process became differentially expressed, indicating that a candidate antibody may be produced in this period. However, it may be too late for these antibodies to overcome Walker-256 cells and to stop the metastasis.

**Figure 7.**
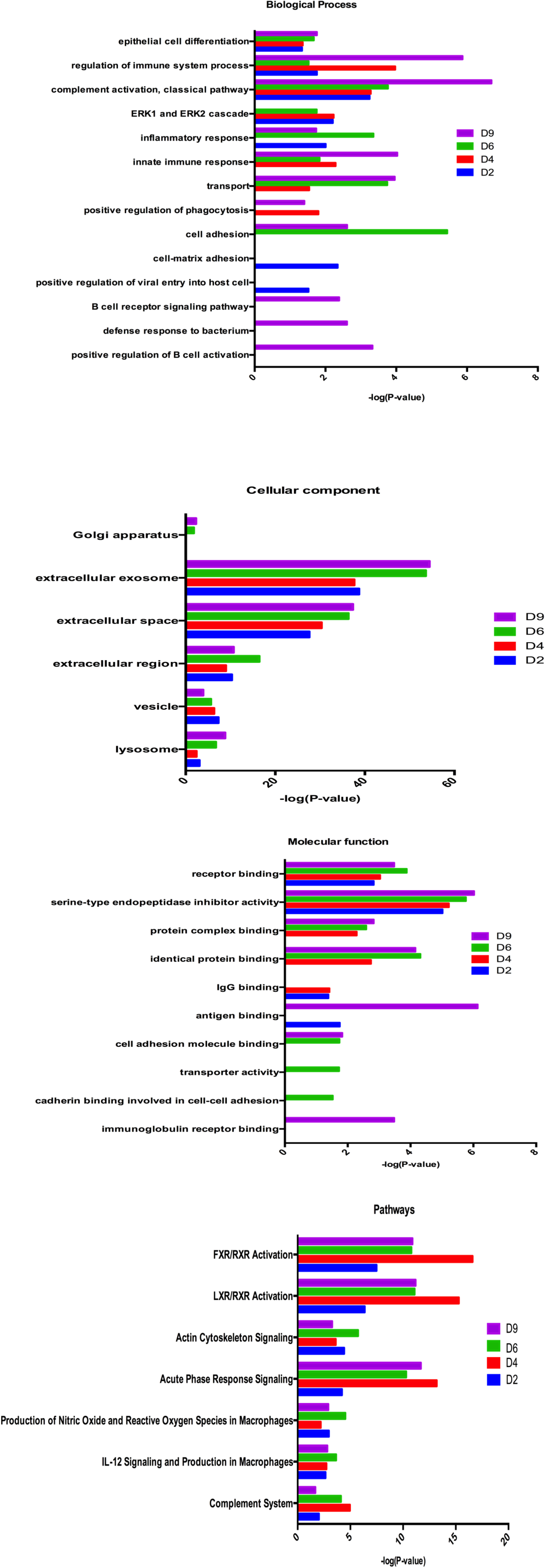
Functional analysis of differential proteins during Walker-256 lung metastatic development. (A) Biological process. (B) Cellular component. (C) Molecular function. (D) Pathways.

The majority of differential proteins in the cellular component category came from extracellular exosomes, the extracellular space, the cellular region, and vesicles. Only a small number of differential proteins were derived from organelles, such as the Golgi apparatus (Figure 7B). This result is consistent with the source of normal urine. In the molecular function category, receptor binding and serine-type endopeptidase inhibitor activity were overrepresented at all time points, while identical protein binding, protein complex binding were overrepresented on days 4, 6, and 9. The transporter activity and cell adhesion molecule binding were both overrepresented on day 6, which is consistent with the cell adhesion and transport process protein differential expression on day 6. On day 9, immunoglobulin receptor binding was overrepresented, but its representation was still consistent with that of the B cell receptor signaling pathway and the positive regulation of B cell activation on day 9 (Figure 7C). It is noteworthy that this molecular function did not appear before day 9.

To identify the major biological pathways involved with the differential urine proteins, we used IPA for canonical pathway enrichment analysis. FXR/RXR activation, LXR/RXR activation, actin cytoskeleton signaling, acute-phase response signaling, IL-12 signaling and production in macrophages, the production of nitric oxide and reactive oxygen species in macrophages, and the complement system were significantly enriched during the whole metastatic progression (Figure 7D). This result indicated that the differential proteins were indeed associated with lung metastatic development.

### Comparison of differential urine proteins in Walker-256 lung carcinoma metastasis rats and Walker-256 subcutaneous rats

There were 15 differential proteins identified specifically when these 62 differential proteins common to 7 lung metastatic rats at one or more of the same time points were compared with Walker-256 subcutaneous model data that our laboratory published before[9]. The comparison procedure is presented in Figure 8. Nine of these 15 differential proteins (NHRF1, CLIC1, EZRI, AMPN, ACY1A, HSP7C, BTD, NID2, and CFAD) were identified at the early stages (day 2 or 4) of lung metastatic development to have homologous human proteins, and their levels continued to change during later lung metastatic stages, suggesting that these proteins have indeed participated in cancer lung metastasis development (Table 2). In addition, eight of these nine differential proteins have been reported to be associated with breast cancer, especially metastasis, while seven of the nine (NHRF1, CLIC1, EZRI, AMPN, ACY1A, HSP7C, and NID2) have been referenced in lung cancer, indicating their roles in the early detection of breast cancer lung metastasis.

**Figure 8.**
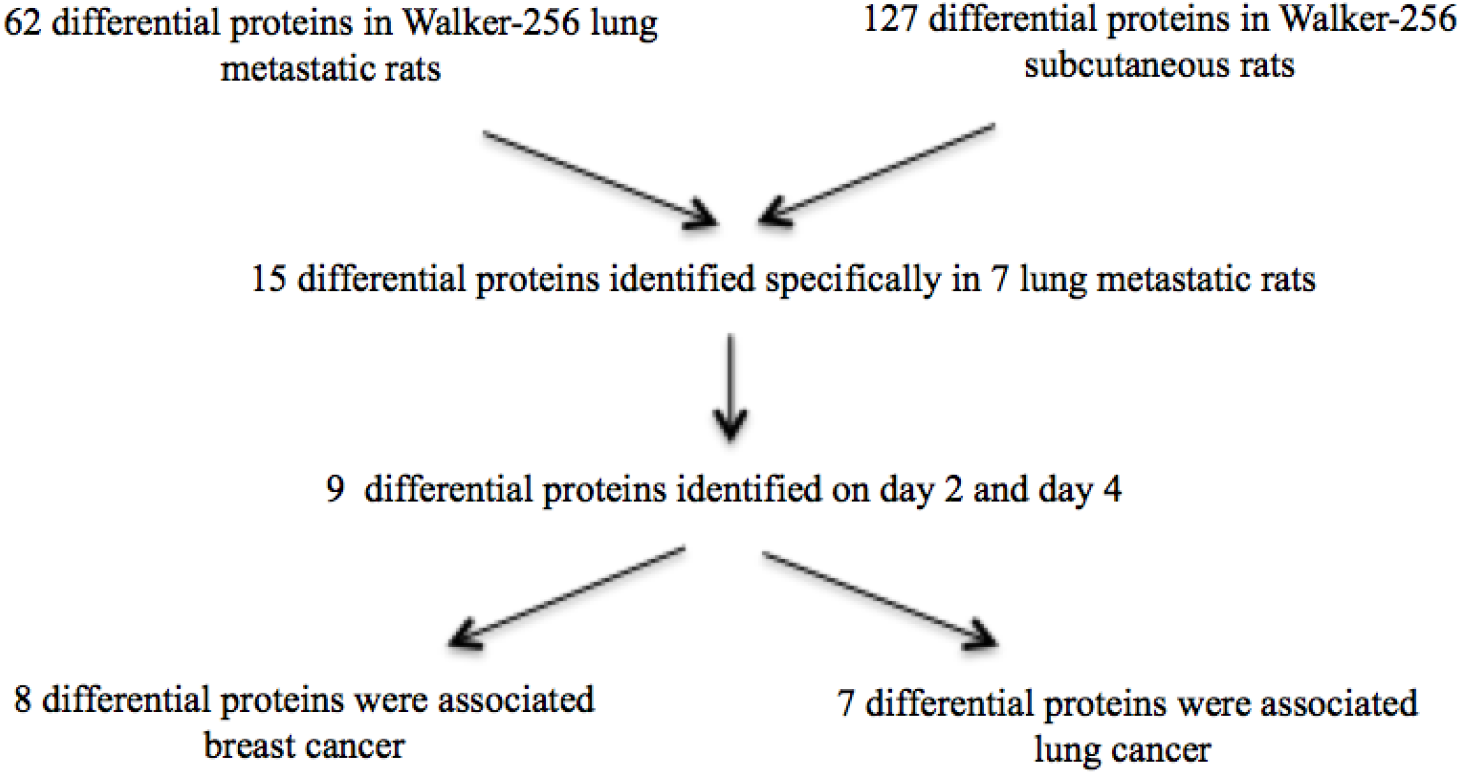
The procedure comparing urine proteins differentially expressed in Walker-256 lung carcinoma metastasis rats and Walker-256 subcutaneous rats.

Seven of these nine differential proteins were associated with both lung cancer and breast cancer, while two of them (BTD and CFAD) were reported in lung or breast cancer. (1) The high expression of Na(+)/H(+) exchange regulatory cofactor NHE-RF1 (NHRF1) is a potential marker of aggressiveness in NSCLC[53]. In addition, the loss of nuclear NHERF1 expression is associated with reduced survival and may serve as a prognostic marker for the routine clinical management of breast cancer patients[54]. Additionally, the expression of NHRF1 can define an immunophenotype of grade 2 invasive breast cancer associated with poor prognosis[57]. (2) Chloride intracellular channel protein 1 (CLIC1) is closely associated with the occurrence and development of lung adenocarcinoma and may thus be used as an effective marker for predicting the prognosis of lung adenocarcinoma[37]. In addition, CLIC1 was also reported to be a potential serological marker for the early detection of breast cancer[38]. (3) Ezrin is an early biomarker for the early diagnosis of lung cancer[33] and a potential prognostic marker of progression in NSCLC[94–96]. In addition, tumor-associated macrophages (TAMs) promote the ezrin phosphorylation-mediated epithelial-mesenchymal transition (EMT) in lung adenocarcinoma through FUT4/LeY-mediated fucosylation[97]. In breast cancer, ezrin regulates focal adhesion and invadopodia dynamics by altering calpain activity to promote breast cancer cell invasion and metastasis[98]. Additionally, ezrin is correlated with cortactin, which facilitates the EMT in breast cancer metastases[35]. (4) The expression of aminopeptidase N (AMPN)/CD13 was reported as a potential unfavorable factor in predicting the efficacy and prognosis of post-operative chemotherapy in NSCLC patients, especially in lung adenocarcinoma patients[66, 67]. Additionally, a high level of circulating AMPN/CD13 was reported to be an independent prognostic factor in patients with NSCLC[69]. In breast cancer patients, the expression of APN/CD13 can serve as a poor prognostic factor in the evaluation of breast cancer prognosis[70]. (5) Aminoacylase-1A is not expressed in SCLC[58]. In addition, the expression level of aminoacylase-1A in human MCF-7 breast cancer cells was altered by 17β-estradiol (E2) treatment and so a might be potent target for treating breast cancer patients[65]. (6) The expression level of cluster of heat shock cognate 71 kDa protein was changed significantly when lung cancer cells were treated with periplocin, which revealed molecular mechanisms underlying the anti-cancer effects of periplocin on lung cancer cells[62]. In MCF-7 and MDA-MB-231 breast cancer cell lines, heat shock cognate 71 kDa protein was reported to be identified as specific phthalic acid-binding proteins [63]. (7) Loss of nidogen-2 significantly promotes lung metastasis of melanoma cells[86]. In human breast cancer specimens, expression of the extracellular protease ADAMTS1 (A disintegrin and metalloprotease with thrombospondin repeats 1) was downregulated, and nidogen proteolysis was partially inhibited, which has implications for vessel integrity[88]. (8) Complement factor D (CFD)/adipsin was overexpressed in BHGc7 cells cultured in conditioned medium, and BHGc7 cells were the first establishment of permanent circulating tumor cell (CTC) lines from blood samples of advanced stage SCLC patients[89]. (9) Biotinidase is a potential serological biomarker for the detection of breast cancer[85].

The presence of some other differential proteins, identified only by Triple TOF 5600^TM^ or Orbitrap Fusion Lumos, cannot be ignored. When the metastasis model data were compared with the Walker-256 subcutaneous data, these differential proteins were screened by the following criteria: (1) only identified at early metastatic stages (days 2 and 4); (2) did not contain the differential proteins annotated in Table 2; (3) all these differential proteins had corresponding homologous proteins; and (4) all these differential proteins exhibited the same trend during the whole lung metastatic periods. The differential lung metastatic proteins only identified in these two mass spectrometers were annotated in Supplement Tables 1 and 2, respectively. There were 17 differential proteins identified by using Triple TOF 5600^TM^, and 13 of these 17 differential proteins have been reported to be associated with lung cancer and breast cancer. Specifically, 10 of them were reported in both lung cancer and breast cancer, while 3 were associated with either lung cancer or breast cancer. There were 24 differential proteins identified by using the Orbitrap Fusion Lumos. Fifteen of these 24 differential proteins are associated with lung cancer and breast cancer. Specifically, 13 of them were reported in both lung cancer and breast cancer, while 2 were associated with either lung cancer or breast cancer. Interestingly, some identified differential proteins were associated with either lung cancer or breast cancer, indicating their novel potential roles as early candidate biomarkers in monitoring Walker-256 lung metastasis progression.

However, in our study, we found that identifying differential proteins by two different mass spectrometers yielded differences. Therefore, we suggest that the use of different mass spectrometers should be considered when conducting clinical applications. Overall, this study was preliminary, and our results indicate that (1) the urine proteome changed significantly, even on the second day after the tail-vein injection of Walker-256 cells and that (2) the urinary differential proteins were different between Walker-256 lung metastatic tumors and Walker-256 subcutaneous tumors. Our results provide a potential possibility to detect early breast cancer lung metastasis, monitor its progression and differentiate it from the same cancer cells grown at other locations.

**Table 2.** Walker-256 lung metastasis differential proteins specifically identified on days 2 and 4.

| Accession | Protein name | ANOVA P |  | Trends | 5600 fold change |  |  |  | Lumos fold change |  |  |  | Lung cancer | Breast cancer |
| --- | --- | --- | --- | --- | --- | --- | --- | --- | --- | --- | --- | --- | --- | --- |
|  |  | 5600 | Lumos |  | D2 | D4 | D6 | D9 | D2 | D4 | D6 | D9 |  |  |
| NHRF1_RAT | Na(+)/H(+) exchange regulatory cofactor<br>NHE-RF1 | 5.08E-06 | 0.006 | ↑ | - | 5.16 | - | 3.66 | 3.97 | 3.91 | 3.23 | 4.91 | [53] | [54, 57] |
| CLIC1_RAT | Chloride intracellular channel protein 1 | 1.56E-10 | 0.12 | ↑ | 7.71 | 11.55 | 6.61 | 10.41 | - | 6.29 | 4.00 | 3.71 | [37] | [38] |
| EZRI_RAT | Ezrin | 6.43E-09 | 0.12 | ↑ | 2.40 | 3.00 | 2.83 | 2.77 | 3.10 | 4.32 | 3.39 | 3.95 | [33, 94-97] | [35, 98] |
| AMPN_RAT | Aminopeptidase N | 1.13E-07 | 0.2 | ↑ | 2.12 | 2.53 | - | 1.53 | - | 1.57 | - | - | [66, 67, 69] | [70] |
| ACY1A_RAT | Aminoacylase-1A | 1.32E-07 | 0.3 | ↑ | 2.66 | 3.55 | 2.38 | 2.45 | - | - | 2.99 | 2.72 | [58] | [65] |
| HSP7C_RAT | Cluster of heat shock cognate 71 kDa protein | 9.29E-09 | 0.63 | ↑ | 2.41 | 2.34 | 1.81 | 2.18 | - | - | 1.72 | 1.67 | [62] | [63] |
| BTD_RAT | Biotinidase | 2.45E-06 | 0.0049 | ↓ |  | 0.60 | 0.53 | 0.49 | 0.56 | 0.53 | - | 0.37 | - | [85] |
| NID2_RAT | Nidogen-2 | 4.09E-08 | 0.038 | ↓ | 0.47 | - | - | - | 0.44 | - | - | - | [86] | [88] |
| CFAD_RAT | Complement factor D | 0.0029 | 0.33 | ↓ | - | 0.60 | 0.59 | - | - | - | 0.64 | - | [89] | - |

**Supplement Table 1.**
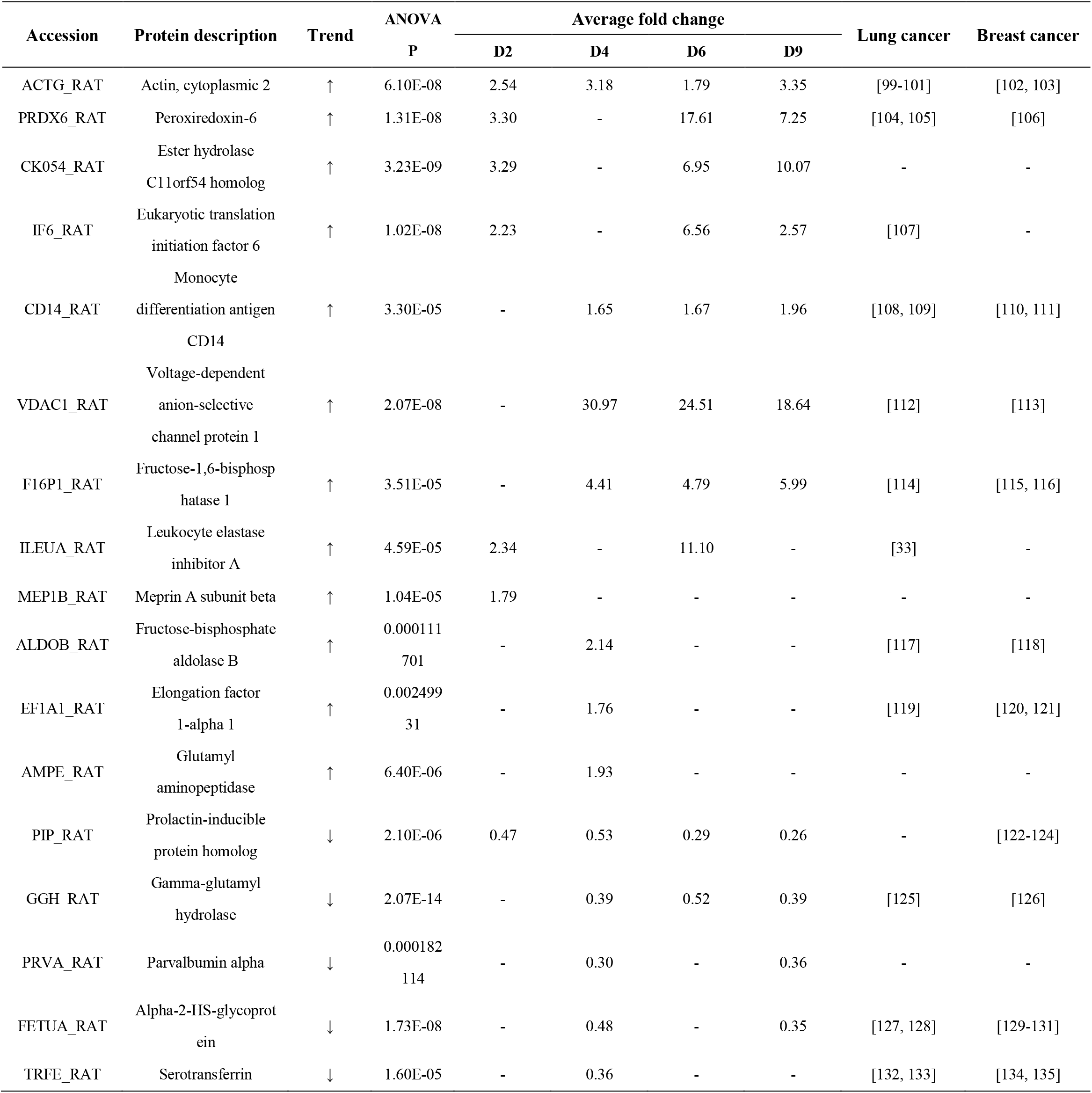
The lung metastasis differential proteins specifically identified with a Triple TOF 5600^TM^.

**Supplement Table 2.** The lung metastasis differential proteins specifically identified with an Orbitrap Fusion Lumos.

| Fusion Lumos |  |  |  |  |  |  |  |  |  |
| --- | --- | --- | --- | --- | --- | --- | --- | --- | --- |
| Accession | Protein description | Trend | ANOVA | Average fold change |  |  |  | Lung cancer | Breast cancer |
|  |  |  | P | D2 | D4 | D6 | D9 |  |  |
| CLIC4_RAT | Chloride intracellular channel protein 4 | ↑ | 0.12 | 23.5 | 20 | 22 | - | [136] | [38, 137] |
| PPIA_RAT | Peptidyl-prolyl cis-trans isomerase A | ↑ | 0.079 | 3.39 | - | 2.46 | 2.87 | [138, 139] | [140, 141] |
| GP2_RAT | Pancreatic secretory granule membrane major glycoprotein GP2 | ↑ | 0.046 | 4.76 | - | 3.18 | 4.66 | - | - |
| NKG2D_RAT | NKG2-D type II integral membrane protein | ↑ | 0.016 | - | 7.89 | 8.33 | 8.11 | - | - |
| ENOA_RAT | Alpha-enolase | ↑ | 0.022 | 1.77 | - | 1.56 | - | [142, 143] | [144] |
| EF1A1_RAT | Elongation factor 1-alpha 1 | ↑ | 0.12 | 3.03 | - | 4.31 | - | [119] | [120, 121] |
| ACTB_RAT | Actin, cytoplasmic 1 | ↑ | 0.27 | 2.30 | - | 2.29 | - | [99, 101, 145] | [102, 103] |
| RHOA_RAT | Transforming protein RhoA | ↑ | 0.26 | 7.25 | - | 5.42 | - | [146, 147] | [148, 149] |
| BASP1_RAT | Brain acid soluble protein 1 | ↑ | 0.22 | 3.33 | - | - | 4.46 | - | - |
| ACY3_RAT | N-acyl-aromatic-L-amino acid amidohydrolase | ↑ | 0.24 | - | 4.30 | - | 3.43 | - | - |
| HS90B_RAT | Heat shock protein HSP 90-beta | ↑ | 0.38 | 8.27 | - | - | - | [150] | [151] |
| CYC_RAT | Cytochrome c, somatic | ↑ | 0.052 | 3.27 | - | - | - | - | - |
| VDAC1_RAT | Voltage-dependent anion-selective channel protein 1 | ↑ | 0.25 | 10.67 | - | - | - | [112] | [113] |
| GDIR1_RAT | Rho GDP-dissociation inhibitor 1 | ↑ | 0.028 | - | 2.92 | - | - | [152] | [153] |
| PRDX1_RAT | Peroxiredoxin-1 | ↑ | 0.62 | - | 3.43 | - | - | [154, 155] | [156] |
| GPC5C_RAT | G-protein coupled receptor family C group 5 member C | ↑ | 0.24 | - | 2.79 | - | - | [157] | - |
| RET4_RAT | Retinol-binding protein 4 | ↓ | 0.0015 | 0.28 | 0.31 | 0.28 | 0.31 | [158] | [159] |
| PPT2_RAT | Lysosomal thioesterase PPT2 | ↓ | 0.088 | 0.59 | 0.62 | 0.36 | - | - | - |
| APOH_RAT | Beta-2-glycoprotein 1 | ↓ | 0.035 | - | 0.39 | 0.45 | 0.44 | [160] | [161, 162] |
| NEUR1_RAT | Sialidase-1 | ↓ | < 0.00010 | 0.60 | - | - | 0.14 | - | - |
| MINP1_RAT | Multiple inositol polyphosphate phosphatase 1 | ↓ | 0.053 | 0.47 | - | - | 0.23 | - | - |
| PCOC1_RAT | Procollagen C-endopeptidase enhancer 1 | ↓ | 0.025 | 0.40 | - | - | 0.43 | - | - |
| HRG_RAT | Histidine-rich glycoprotein | ↓ | 0.42 | - | 0.33 | 0.42 | - | - | [163] |

## Acknowledgements

This research was supported by the National Key Research and Development Program of China (2016 YFC 1306300), Key Basic Research Program of the Ministry of Science and Technology of China (2013FY114100), Beijing Natural Science Foundation (7173264, 7172076), Beijing cooperative construction project (110651103), Beijing Normal University (11100704), Peking Union Medical College Hospital (2016-2.27). The funders had no role in study design, data collection and analysis, decision to publish, or preparation of the manuscript.

## Conflict of Interest/Disclosure Statement

The authors have no conflict of interest to report.

## References

1. Nguyen DX, Bos PD, Massague J. Metastasis: from dissemination to organ-specific colonization. Nat Rev Cancer. 2009;9(4):274–84. doi: 10.1038/nrc2622. PubMed PMID: 19308067.

2. Guan X. Cancer metastases: challenges and opportunities. Acta Pharm Sin B. 2015;5(5):402–18. doi: 10.1016/j.apsb.2015.07.005. PubMed PMID: 26579471; PubMed Central PMCID: PMCPMC4629446.

3. Monica J. Lewis JL, Emily Falk Libby, Minnkyong Lee, Nigel P.S., Crawford DRH. SIN3A and SIN3B differentially regulate breast cancer metastasis. Oncotarget.2016;8(48):78713–25.

4. Clever D, Roychoudhuri R, Constantinides MG, Askenase MH, Sukumar M, Klebanoff CA, et al. Oxygen Sensing by T Cells Establishes an Immunologically Tolerant Metastatic Niche. Cell. 2016;166(5):1117–31 e14. doi: 10.1016/j.cell.2016.07.032. PubMed PMID: 27565342; PubMed Central PMCID: PMCPMC5548538.

5. Gao Y Are Urinary Biomarkers from Clinical Studies Biomarkers of Disease or Biomarkers of Medicine? MOJ Proteomics & Bioinformatics. 2014;1(5). doi: 10.15406/mojpb.2014.01.00028.

6. Gao Y. Urine-an untapped goldmine for biomarker discovery? Science China Life sciences. 2013;56(12):1145–6. doi: 10.1007/s11427-013-4574-1. PubMed PMID: 24271956.

7. Wu J, Gao Y Physiological conditions can be reflected in human urine proteome and metabolome. Expert review of proteomics. 2015;12(6):623–36. doi: 10.1586/14789450.2015.1094380. PubMed PMID: 26472227.

8. Gao Y. Roadmap to the Urine Biomarker Era. MOJ Proteomics & Bioinformatics. 2014;1(1):00005.

9. Wu J, Guo Z, Gao Y Dynamic changes of urine proteome in a Walker 256 tumor-bearing rat model. Cancer Med. 2017. doi: 10.1002/cam4.1225. PubMed PMID: 28980450.

10. Beretov J, Wasinger VC, Millar EK, Schwartz P, Graham PH, Li Y. Proteomic Analysis of Urine to Identify Breast Cancer Biomarker Candidates Using a Label-Free LC-MS/MS Approach. PLoS One. 2015;10(11):e0141876. doi: 10.1371/journal.pone.0141876. PubMed PMID: 26544852; PubMed Central PMCID: PMCPMC4636393.

11. Beretov J, Wasinger VC, Graham PH, Millar EK, Kearsley JH, Li Y. Proteomics for Breast Cancer Urine Biomarkers. 2014;63:123–67. doi: 10.1016/b978-0-12-800094-6.00004-2.

12. Changsheng Dong M, RuiXin Wu, MD, Jing Wu, PhD, Jing Guo, MD, Fangyuan Wang, MS, Yanli Fu, PhD, Qing Wang, MS, Ling Xu, Professor, and Juyong Wang, Professor. Evaluation of Bone Cancer Pain Induced by Different Doses of Walker 256 Mammary Gland Carcinoma Cells. Pain Physician. 2016;19:E1063–E77.

13. FreireGarabal M, NunezIglesias, M., Losada, C., PereiroRaposo, M., CastroBolano, C., Heras, J., Riveiro, P., Mayan, J., ReyMendez, M. Stimulatory effects of amphetamine on the development of Walker-256 carcinoma lung metastases in rats. Oncology Reports. 1996;3(3):201–4.

14. Wisniewski JR, Zougman A, Nagaraj N, Mann M. Universal sample preparation method for proteome analysis. Nat Methods. 2009;6(5):359–62. doi: 10.1038/nmeth.1322. PubMed PMID: 19377485.

15. Huang da W, Sherman BT, Lempicki RA. Systematic and integrative analysis of large gene lists using DAVID bioinformatics resources. Nat Protoc. 2009;4(1):44–57. doi: 10.1038/nprot.2008.211. PubMed PMID: 19131956.

16. Fukuda T, Nomura M, Kato Y, Tojo H, Fujii K, Nagao T, et al. A selected reaction monitoring mass spectrometric assessment of biomarker candidates diagnosing large-cell neuroendocrine lung carcinoma by the scaling method using endogenous references. PLoS One. 2017;12(4):e0176219. doi: 10.1371/journal.pone.0176219. PubMed PMID: 28448532; PubMed Central PMCID: PMCPMC5407814.

17. Tzu-Wen Lin H-TC, Chein-Hung Chen, Chung-Hsuan Chen, Sheng-Wei Lin, Tsui-Ling Hsu and Chi-Huey Wong. Galectin-3 binding protein and galectin-1 interaction in breast cancer cell aggregation and metastasis. Journal of the American Chemical Society. 2015;137(30):9685–93. PubMed PMID: 26168351.

18. Beibei Sun WG, Song Hu, Feng Yao, Keke Yu, Jie Xing, Ronghua Wang, Hongyong Song, Yueling Liao, Tong Wang, Pengfei Jiang, Baohui Han, Jiong Deng. Gprc5a-knockout mouse lung epithelial cells predicts ceruloplasmin, lipocalin 2 and periostin as potential biomarkers at early stages of lung tumorigenesis. Oncotarget. 2017;8(8):13532–44. PubMed PMID: 28088789

19. Ören B,Urosevic J, Mertens C, Mora J, Guiu M, Gomis RR, et al. Tumour stroma-derived lipocalin-2 promotes breast cancer metastasis. The Journal of Pathology. 2016;239(3):274–85. doi: 10.1002/path.4724. PubMed PMID: 27038000

20. Kotteas EA, Gkiozos I, Tsagkouli S, Bastas A, Ntanos I, Saif MW, et al. Soluble ICAM-1 levels in small-cell lung cancer: prognostic value for survival and predictive significance for response during chemotherapy. Med Oncol. 2013;30(3):662. doi: 10.1007/s12032-013-0662-0. PubMed PMID: 23884579.

21. Schroder C, Witzel I, Muller V, Krenkel S, Wirtz RM, Janicke F, et al. Prognostic value of intercellular adhesion molecule (ICAM)-1 expression in breast cancer. J Cancer Res Clin Oncol. 2011;137(8):1193–201. doi: 10.1007/s00432-011-0984-2. PubMed PMID: 21590495.

22. Fiala O, Hosek P, Pesek M, Finek J, Racek J, Buchler T, et al. Prognostic role of serum C-reactive protein in patients with advanced-stage NSCLC treated with pemetrexed. Neoplasma. 2017;64(4):605–10. doi: 10.4149/neo_2017_416. PubMed PMID: 28485168.

23. Frydenberg H, Thune I, Lofterod T, Mortensen ES, Eggen AE, Risberg T, et al. Pre-diagnostic high-sensitive C-reactive protein and breast cancer risk, recurrence, and survival. Breast Cancer Res Treat. 2016;155(2):345–54. doi: 10.1007/s10549-015-3671-1. PubMed PMID: 26740213.

24. Wang Y, Chen Z, Chen J, Pan J, Zhang W, Pan Q, et al. The diagnostic value of apolipoprotein E in malignant pleural effusion associated with non-small cell lung cancer. Clin Chim Acta. 2013;421:230–5. doi: 10.1016/j.cca.2013.03.013. PubMed PMID: 23523589.

25. Su WP, Chen YT, Lai WW, Lin CC, Yan JJ, Su WC. Apolipoprotein E expression promotes lung adenocarcinoma proliferation and migration and as a potential survival marker in lung cancer. Lung Cancer. 2011;71(1):28–33. doi: 10.1016/j.lungcan.2010.04.009. PubMed PMID: 20430468.

26. Xu X, Wan J, Yuan L, Ba J, Feng P, Long W, et al. Serum levels of apolipoprotein E correlates with disease progression and poor prognosis in breast cancer. Tumour Biol. 2016. doi: 10.1007/s13277-016-5453-8. PubMed PMID: 27709551.

27. Szoke T, Kayser K, Baumhakel JD, Trojan I, Furak J, Tiszlavicz L, et al. Prognostic significance of endogenous adhesion/growth-regulatory lectins in lung cancer. Oncology. 2005;69(2):167–74. doi: 10.1159/000087841. PubMed PMID: 16127288.

28. Srinivasan N, Bane SM, Ahire SD, Ingle AD, Kalraiya RD. Poly N-acetyllactosamine substitutions on N- and not O-oligosaccharides or Thomsen-Friedenreich antigen facilitate lung specific metastasis of melanoma cells via galectin-3. Glycoconj J. 2009;26(4):445–56. doi: 10.1007/s10719-008-9194-9. PubMed PMID: 18949555.

29. Noma N, Simizu S, Kambayashi Y, Kabe Y, Suematsu M, Umezawa K. Involvement of NF-kappaB-mediated expression of galectin-3-binding protein in TNF-alpha-induced breast cancer cell adhesion. Oncol Rep. 2012;27(6):2080–4. doi: 10.3892/or.2012.1733. PubMed PMID: 22447108.

30. White MJ, Roife D, Gomer RH. Galectin-3 Binding Protein Secreted by Breast Cancer Cells Inhibits Monocyte-Derived Fibrocyte Differentiation. J Immunol. 2015;195(4):1858–67. doi: 10.4049/jimmunol.1500365. PubMed PMID: 26136428; PubMed Central PMCID: PMCPMC4530092.

31. Li Q, Gao H, Xu H, Wang X, Pan Y, Hao F, et al. Expression of ezrin correlates with malignant phenotype of lung cancer, and in vitro knockdown of ezrin reverses the aggressive biological behavior of lung cancer cells. Tumour Biol. 2012;33(5):1493–504. doi: 10.1007/s13277-012-0400-9. PubMed PMID: 22528947.

32. Aman Wang CL, Zhen Ning2, Wei Gao, Yunpeng Xie, Ningning Zhang, Jinxiao Liang, Faisal S. Abbasi, Qiu Yan, Jiwei Liu. Tumor-associated macrophages promote Ezrin phosphorylation-mediated epithelial-mesenchymal transition in lung adenocarcinoma through FUT4/LeY up-regulation. Oncotarget. 2017;8(17):28247–59. PubMed PMID: 28423676

33. Pastor MD, Nogal A, Molina-Pinelo S, Melendez R, Salinas A, Gonzalez De la Pena M, et al. Identification of proteomic signatures associated with lung cancer and COPD. J Proteomics. 2013;89:227–37. doi: 10.1016/jjprot.2013.04.037. PubMed PMID: 23665002.

34. Murni H. Jais MU, Reena R. Md Zin, PhD,Nur A. Muhd Hanapi, BBiomed, and Siti A. Md Ali, DCP. Ezrin is Significantly Overexpressed in Luminal A, Luminal B, and HER2 Subtype Breast Cancer. Appl Immunohistochem Mol Morphol 2017;25(1):44–8. PubMed PMID: 26469327.

35. He J, Ma G, Qian J, Zhu Y, Liang M, Yao N, et al. Interaction Between Ezrin and Cortactin in Promoting Epithelial to Mesenchymal Transition in Breast Cancer Cells. Medical Science Monitor. 2017;23:1583–96. doi: 10.12659/msm.904124. PubMed PMID: 28364518

36. Fentiman IS, Allen DS. gamma-Glutamyl transferase and breast cancer risk. Br J Cancer. 2010;103(1):90–3. doi: 10.1038/sj.bjc.6605719. PubMed PMID: 20517309; PubMed Central PMCID: PMCPMC2905293.

37. Wang W, Xu X, Wang W, Shao W, Li L, Yin W, et al. The expression and clinical significance of CLIC1 and HSP27 in lung adenocarcinoma. Tumour Biol. 2011;32(6):1199–208. doi: 10.1007/s13277-011-0223-0. PubMed PMID: 21858536.

38. Gromov P, Gromova I, Bunkenborg J, Cabezon T, Moreira JM, Timmermans-Wielenga V, et al. Up-regulated proteins in the fluid bathing the tumour cell microenvironment as potential serological markers for early detection of cancer of the breast. Mol Oncol. 2010;4(1):65–89. doi: 10.1016/j.molonc.2009.11.003. PubMed PMID: 20005186; PubMed Central PMCID: PMCPMC5527961.

39. Masahide Tokunou TN, Yukihito Saitoh, Hiroji Imamura, Michiie Sakamoto, and Setsuo Hirohashi. Altered Expression of the ERM Proteins in Lung Adenocarcinoma. Lab Invest. 2000;80(11):1643–50. PubMed PMID: 11092524.

40. Bartova M, Hlavaty J, Tan Y, Singer C, Pohlodek K, Luha J, et al. Expression of ezrin and moesin in primary breast carcinoma and matched lymph node metastases. Clin Exp Metastasis. 2017;34(5):333–44. doi: 10.1007/s10585-017-9853-y. PubMed PMID: 28624994.

41. Charafe-Jauffret E, Monville F, Bertucci F, Esterni B, Ginestier C, Finetti P, et al. Moesin expression is a marker of basal breast carcinomas. Int J Cancer. 2007;121(8):1779–85. doi: 10.1002/ijc.22923. PubMed PMID: 17594689.

42. Hong H, Yu H, Yuan J, Guo C, Cao H, Li W, et al. MicroRNA-200b Impacts Breast Cancer Cell Migration and Invasion by Regulating Ezrin-Radixin-Moesin. Medical Science Monitor. 2016;22:1946–52. doi: 10.12659/msm.896551. PubMed PMID: 27276064.

43. Hmmier A, O'Brien ME, Lynch V, Clynes M, Morgan R, Dowling P. Proteomic analysis of bronchoalveolar lavage fluid (BALF) from lung cancer patients using label-free mass spectrometry. BBA Clin. 2017;7:97–104. doi: 10.1016/j.bbacli.2017.03.001. PubMed PMID: 28331811; PubMed Central PMCID: PMCPMC5357681.

44. Ajona D, Pajares MJ, Corrales L, Perez-Gracia JL, Agorreta J, Lozano MD, et al. Investigation of complement activation product c4d as a diagnostic and prognostic biomarker for lung cancer. J Natl Cancer Inst. 2013;105(18):1385–93. doi: 10.1093/jnci/djt205. PubMed PMID: 23940286; PubMed Central PMCID: PMCPMC3776260.

45. Ortea I, Rodriguez-Ariza A, Chicano-Galvez E, Arenas Vacas MS, Jurado Gamez B. Discovery of potential protein biomarkers of lung adenocarcinoma in bronchoalveolar lavage fluid by SWATH MS data-independent acquisition and targeted data extraction. J Proteomics. 2016;138:106–14. doi: 10.1016/jjprot.2016.02.010. PubMed PMID: 26917472.

46. Ajona D, Razquin C, Pastor MD, Pajares MJ, Garcia J, Cardenal F, et al. Elevated levels of the complement activation product C4d in bronchial fluids for the diagnosis of lung cancer. PLoS One. 2015;10(3):e0119878. doi: 10.1371/journal.pone.0119878. PubMed PMID: 25799154; PubMed Central PMCID: PMCPMC4370816.

47. Michlmayr A, Bachleitner-Hofmann T, Baumann S, Marchetti-Deschmann M, Rech-Weichselbraun I, Burghuber C, et al. Modulation of plasma complement by the initial dose of epirubicin/docetaxel therapy in breast cancer and its predictive value. Br J Cancer. 2010;103(8):1201–8. doi: 10.1038/sj.bjc.6605909. PubMed PMID: 20877360; PubMed Central PMCID: PMCPMC2967072.

48. Ayyub A, Saleem M, Fatima I, Tariq A, Hashmi N, Musharraf SG. Glycosylated Alpha-1-acid glycoprotein 1 as a potential lung cancer serum biomarker. Int J Biochem Cell Biol. 2016;70:68–75. doi: 10.1016/j.biocel.2015.11.006. PubMed PMID: 26563422.

49. Ganz PA BM, Ma PY, Elashoff RM. Monitoring the Therapy of Lung Cancer with a-1-Acid Glycoprotein1. Cancer Res. 1984;44(11):5415–21. PubMed PMID: 6435869.

50. René Bruno RO, Jocelyne Berille, Philip Chaikin, Nicole Vivier, Luz Hammershaimb,Gerald R. Rhodes, and James R. Rigas. Alpha-1-Acid Glycoprotein As an Independent Predictor for Treatment Effects and a Prognostic Factor of Survival in Patients with Non-small Cell Lung Cancer Treated with Docetaxel. Clin Cancer Res 2003;9(3):1077–82. PubMed PMID: 12631610.

51. J. G. ROBERTS JWKAMB. Serum a-I-acid glycoprotein as an index of dissemination in-breast cancer. Br J Surg. 1975;62(10):816–9. PubMed PMID: 1191942.

52. Stephen HM, Khoury RJ, Majmudar PR, Blaylock T, Hawkins K, Salama MS, et al. Epigenetic suppression of neprilysin regulates breast cancer invasion. Oncogenesis. 2016;5:e207. doi: 10.1038/oncsis.2016.16. PubMed PMID: 26950599; PubMed Central PMCID: PMCPMC4815048.

53. Mangia A, Partipilo G, Schirosi L, Saponaro C, Galetta D, Catino A, et al. Fine Needle Aspiration Cytology: A Tool to Study NHERF1 Expression as a Potential Marker of Aggressiveness in Lung Cancer. Mol Biotechnol. 2015;57(6):549–57. doi: 10.1007/s12033-015-9848-3. PubMed PMID: 25744438.

54. Paradiso A, Scarpi E, Malfettone A, Addati T, Giotta F, Simone G, et al. Nuclear NHERF1 expression as a prognostic marker in breast cancer. Cell Death Dis. 2013;4:e904. doi: 10.1038/cddis.2013.439. PubMed PMID: 24201803; PubMed Central PMCID: PMCPMC3847317.

55. Georgescu MM. NHERF1: molecular brake on the PI3K pathway in breast cancer. Breast Cancer Res. 2008;10(2):106. doi: 10.1186/bcr1992. PubMed PMID: 18430260; PubMed Central PMCID: PMCPMC2397532.

56. Karn T, Ruckhaberle E, Hanker L, Muller V, Schmidt M, Solbach C, et al. Gene expression profiling of luminal B breast cancers reveals NHERF1 as a new marker of endocrine resistance. Breast Cancer Res Treat. 2011;130(2):409–20. doi: 10.1007/s10549-010-1333-x. PubMed PMID: 21203899.

57. Andrea Malfettone CS, Angelo Paradiso, Giovanni Simone3 and Annita Mangia. Peritumoral vascular invasion and NHERF1 expression define an immunophenotype of grade invasive breast cancer associated with poor prognosis. BMC Cancer. 2012;12:106. PubMed PMID: 22439624.

58. York E. Miller JDM, and Adi F. Gazdar. Lack of Expression of Aminoacylase-1 in Small Cell Lung Cancer. Evidence for Inactivation of Genes Encoded by Chromosome 3p. TheJournalofClinicalInvestigation,Inc. 1989;83:2120–4. PubMed PMID: 2542383.

59. Ocak S, Pedchenko TV, Chen H, Harris FT, Qian J, Polosukhin V, et al. Loss of polymeric immunoglobulin receptor expression is associated with lung tumourigenesis. Eur Respir J. 2012;39(5):1171–80. doi: 10.1183/09031936.00184410. PubMed PMID: 21965228; PubMed Central PMCID: PMCPMC3717253.

60. Ting Xiao WY, Lei Li,Zhi Hu,Ying Ma, Liyan Jiao,Jinfang Ma, Yun Cai, Dongmei Lin, Suping Guo, Naijun Han,Xuebing Di, Min Li, Dechao Zhang KS, e Jinsong Yuan, Hongwei Zheng, Meixia Gao, Jie He, Susheng Shi, Wuju Li, Ningzhi Xu, Husheng Zhang, Yan Liu, Kaitai Zhang, Yanning Gao, Xiaohong Qian and Shujun Cheng. An Approach to Studying Lung Cancer-related Proteins in Human Blood. Mol Cell Proteomics. 2005;4(10):1480–6. doi: 10.1074/. PubMed PMID: 15970581.

61. Prasad A, Fernandis AZ, Rao Y, Ganju RK. Slit Protein-mediated Inhibition of CXCR4-induced Chemotactic and Chemoinvasive Signaling Pathways in Breast Cancer Cells. Journal of Biological Chemistry. 2004;279(10):9115–24. doi: 10.1074/jbc.M308083200. PubMed PMID: 14645233.

62. Lu Z, Song Q, Yang J, Zhao X, Zhang X, Yang P, et al. Comparative proteomic analysis of anti-cancer mechanism by periplocin treatment in lung cancer cells. Cell Physiol Biochem. 2014;33(3):859–68. doi: 10.1159/000358658. PubMed PMID: 24685647.

63. Liang SS, Wang TN, Tsai EM. Analysis of protein-protein interactions in MCF-7 and MDA-MB-231 cell lines using phthalic acid chemical probes. Int J Mol Sci. 2014;15(11):20770–88. doi: 10.3390/ijms151120770. PubMed PMID: 25402641; PubMed Central PMCID: PMCPMC4264195.

64. ELIAS A. KOTTEAS PB, IOANNIS GKIOZOS, SOFIA TSAGKOULI, GEORGE TSOUKALAS and KONSTANTINOS N. SYRIGOS. The Intercellular Cell Adhesion Molecule-1 (ICAM-1) in Lung Cancer: Implications for Disease Progression and Prognosis. Anticancer Res. 2014;34(9):4665–72. PubMed PMID: 25202042.

65. Lee SU, Kim BT, Min YK, Kim SH. Protein profiling and transcript expression levels of heat shock proteins in 17beta-estradiol-treated human MCF-7 breast cancer cells. Cell Biol Int. 2006;30(12):983–91. doi: 10.1016/j.cellbi.2006.07.005. PubMed PMID: 16962797.

66. Zhang Q, Wang J, Zhang H, Zhao D, Zhang Z, Zhang S. Expression and clinical significance of aminopeptidase N/CD13 in non-small cell lung cancer. J Cancer Res Ther. 2015;11(1):223–8. doi: 10.4103/0973-1482.138007. PubMed PMID: 25879366.

67. Tokuhara T, Hattori N, Ishida H, Hirai T, Higashiyama M, Kodama K, et al. Clinical significance of aminopeptidase N in non-small cell lung cancer. Clin Cancer Res. 2006;12(13):3971–8. doi: 10.1158/1078-0432.CCR-06-0338. PubMed PMID: 16818694.

68. Ito S, Miyahara R, Takahashi R, Nagai S, Takenaka K, Wada H, et al. Stromal aminopeptidase N expression: correlation with angiogenesis in non-small-cell lung cancer. Gen Thorac Cardiovasc Surg. 2009;57(11):591–8. doi: 10.1007/s11748-009-0445-x. PubMed PMID: 19908113.

69. Murakami H, Yokoyama A, Kondo K, Nakanishi S, Kohno N, Miyake M. Circulating aminopeptidase N/CD13 is an independent prognostic factor in patients with non-small cell lung cancer. Clin Cancer Res. 2005;11(24 Pt 1):8674–9. doi: 10.1158/1078-0432.CCR-05-1005. PubMed PMID: 16361553.

70. Ranogajec I, Jakic-Razumovic J, Puzovic V, Gabrilovac J. Prognostic value of matrix metalloproteinase-2 (MMP-2), matrix metalloproteinase-9 (MMP-9) and aminopeptidase N/CD13 in breast cancer patients. Med Oncol. 2012;29(2):561–9. doi: 10.1007/s12032-011-9984-y. PubMed PMID: 21611838.

71. Hou L, Zhao X, Wang P, Ning Q, Meng M, Liu C. Antitumor activity of antimicrobial peptides containing CisoDGRC in CD13 negative breast cancer cells. PLoS One. 2013;8(1):e53491. doi: 10.1371/journal.pone.0053491. PubMed PMID: 23326440; PubMed Central PMCID: PMCPMC3543424.

72. Orecchioni S, Gregato G, Martin-Padura I, Reggiani F, Braidotti P, Mancuso P, et al. Complementary populations of human adipose CD34+ progenitor cells promote growth, angiogenesis, and metastasis of breast cancer. Cancer Res. 2013;73(19):5880–91. doi: 10.1158/0008-5472.CAN-13-0821. PubMed PMID: 23918796.

73. Fan J, Yu H, Lv Y, Yin L. Diagnostic and prognostic value of serum thioredoxin and DJ-1 in non-small cell lung carcinoma patients. Tumour Biol. 2016;37(2):1949–58. doi: 10.1007/s13277-015-3994-x. PubMed PMID: 26334622.

74. Park BJ CM, Kim IH. Thioredoxin 1 as a serum marker for breast cancer and its use in combination with CEA or CA15-3 for improving the sensitivity of breast cancer diagnoses. BMC Res Notes. 2014;6:7:. PubMed PMID: 24393391.

75. Ada TG, Ada AO, Kunak SC, Alpar S, Gulhan M, Iscan M. Association between glutathione S-transferase omega 1 A140D polymorphism in the Turkish population and susceptibility to non-small cell lung cancer. Arh Hig Rada Toksikol. 2013;64(2):61–7. doi: 10.2478/10004-1254-64-2013-2302. PubMed PMID: 23819933.

76. Lu H, Chen I, Shimoda LA, Park Y, Zhang C, Tran L, et al. Chemotherapy-Induced Ca(2+) Release Stimulates Breast Cancer Stem Cell Enrichment. Cell Rep. 2017;18(8):1946–57. doi: 10.1016/j.celrep.2017.02.001. PubMed PMID: 28228260.

77. Zeng Q, Xue N, Dai D, Xing S, He X, Li S, et al. A Nomogram based on Inflammatory Factors C-Reactive Protein and Fibrinogen to Predict the Prognostic Value in Patients with Resected Non-Small Cell Lung Cancer. J Cancer. 2017;8(5):744–53. doi: 10.7150/jca.17423. PubMed PMID: 28382136; PubMed Central PMCID: PMCPMC5381162.

78. Luo J, Song J, Feng P, Wang Y, Long W, Liu M, et al. Elevated serum apolipoprotein E is associated with metastasis and poor prognosis of non-small cell lung cancer. Tumour Biol. 2016;37(8):10715–21. doi: 10.1007/s13277-016-4975-4. PubMed PMID: 26873483.

79. Anic GM, Weinstein SJ, Mondul AM, Mannisto S, Albanes D. Serum vitamin D, vitamin D binding protein, and lung cancer survival. Lung Cancer. 2014;86(3):297–303. doi: 10.1016/j.lungcan.2014.10.008. PubMed PMID: 25456734; PubMed Central PMCID: PMCPMC4267905.

80. Turner AM, McGowan L, Millen A, Rajesh P, Webster C, Langman G, et al. Circulating DBP level and prognosis in operated lung cancer: an exploration of pathophysiology. Eur Respir J. 2013;41(2):410–6. doi: 10.1183/09031936.00002912. PubMed PMID: 22556021.

81. Thyer L, Ward E, Smith R, Fiore MG, Magherini S, Branca JJ, et al. A novel role for a major component of the vitamin D axis: vitamin D binding protein-derived macrophage activating factor induces human breast cancer cell apoptosis through stimulation of macrophages. Nutrients. 2013;5(7):2577–89. doi: 10.3390/nu5072577. PubMed PMID: 23857228; PubMed Central PMCID: PMCPMC3738989.

82. Fanrong Zhang LY, Jiaoyue Jin, Kaiyan Chen, Nan Zhang, Junzhou Wu, Yimin Zhang and Dan Su. The C-reactive protein/albumin ratio predicts long-term outcomes of patients with operable non-small cell lung cancer. Oncotarget. 2017;8(5):8835–42. PubMed PMID: 27823974

83. Yao X, Wang X, Han H, Duan Q, Khan U, Hu Y. Changes of serum albumin level and systemic inflammatory response in inoperable non-small cell lung cancer patients after chemotherapy. Journal of Cancer Research and Therapeutics. 2014;10(4):1019. doi: 10.4103/0973-1482.137953.

84. Liu L, Bi Y, Zhou M, Chen X, He X, Zhang Y, et al. Biomimetic Human Serum Albumin Nanoparticle for Efficiently Targeting Therapy to Metastatic Breast Cancers. ACS Appl Mater Interfaces. 2017;9(8):7424–35. doi: 10.1021/acsami.6b14390. PubMed PMID: 28150932.

85. Kang U-B, Ahn Y, Lee JW, Kim Y-H, Kim J, Yu M-H, et al. Differential profiling of breast cancer plasma proteome by isotope-coded affinity tagging method reveals biotinidase as a breast cancer biomarker. BMC Cancer. 2010;10(1). doi: 10.1186/1471-2407-10-114. PubMed PMID: 20346108.

86. Mokkapati S, Bechtel M, Reibetanz M, Miosge N, Nischt R. Absence of the basement membrane component nidogen 2, but not of nidogen 1, results in increased lung metastasis in mice. J Histochem Cytochem. 2012;60(4):280–9. doi: 10.1369/0022155412436586. PubMed PMID: 22260998; PubMed Central PMCID: PMCPMC3351237.

87. Annie Wai Yeeng Chai AKLC, Wei Dai, Josephine Mun Yee Ko, Joseph Chok Yan Ip, Kwok Wah Chan, Dora Lai-Wan Kwong, Wai Tong Ng, Anne Wing Mui Lee, Roger Kai Cheong Ngan, Chun Chung Yau, Stewart Yuk Tung, Victor Ho Fun Lee, Alfred King-Yin Lam, Suja Pillai, Simon Law, Maria Li Lung. Metastasis-suppressing NID2, an epigenetically-silenced gene, in the pathogenesis of nasopharyngeal carcinoma and esophageal squamous cell carcinoma. Oncotarget. 2016;7(48):78859–71. PubMed PMID: 27793011.

88. Martino-Echarri E, Fernandez-Rodriguez R, Rodriguez-Baena FJ, Barrientos-Duran A, Torres-Collado AX, Plaza-Calonge Mdel C, et al. Contribution of ADAMTS1 as a tumor suppressor gene in human breast carcinoma. Linking its tumor inhibitory properties to its proteolytic activity on nidogen-1 and nidogen-2. Int J Cancer. 2013;133(10):2315–24. doi: 10.1002/ijc.28271. PubMed PMID: 23681936.

89. Hamilton G, Rath B, Klameth L, Hochmair MJ. Small cell lung cancer: Recruitment of macrophages by circulating tumor cells. Oncoimmunology. 2016;5(3):e1093277. doi: 10.1080/2162402X.2015.1093277. PubMed PMID: 27141354; PubMed Central PMCID: PMCPMC4839345.

90. Okano T, Seike M, Kuribayashi H, Soeno C, Ishii T, Kida K, et al. Identification of haptoglobin peptide as a novel serum biomarker for lung squamous cell carcinoma by serum proteome and peptidome profiling. Int J Oncol. 2016;48(3):945–52. doi: 10.3892/ijo.2016.3330. PubMed PMID: 26783151; PubMed Central PMCID: PMCPMC4750543.

91. Custodio A, Lopez-Farre AJ, Zamorano-Leon JJ, Mateos-Caceres PJ, Macaya C, Caldes T, et al. Changes in the expression of plasma proteins associated with thrombosis in BRCA1 mutation carriers. J Cancer Res Clin Oncol. 2012;138(5):867–75. doi: 10.1007/s00432-012-1161-y. PubMed PMID: 22311183.

92. Jar-Yi Ho R-JH, Chih-Hsi Wu, Guo-Shiou Liao, Hong-Wei Gao, Tong-Hong Wang, Cheng-Ping Yu. Reduced miR-550a-3p leads to breast cancer initiation, growth, and metastasis by increasing levels of ERK1 and 2. Oncotarget. 2016;7(33):53853–68. PubMed PMID: 27462780

93. FlorinNiculescu HR, Maria Retegan, and Roman Vlaicu. Persistent Complement Activation on Tumor Cells in Breast Cancer. Am J Pathol 1992;140(5):1039–43. PubMed PMID: 1374587

94. Zhang XQ, Chen GP, Wu T, Yan JP, Zhou JY. Expression and clinical significance of ezrin in non–small-cell lung cancer. Clin Lung Cancer. 2012;13(3):196–204. doi: 10.1016/j.cllc.2011.04.002. PubMed PMID: 22137559.

95. Tiefeng Jin JJ, Xiangyu Li, Songnan Zhang, Yun Ho Choi, Yingshi Piao, Xionghu Shen3 and Zhenhua Lin. Prognostic implications of ezrin and phosphorylated ezrin expression in non-small cell lung cancer. BMC Cancer. 2014;14:191.

96. Lee HW, Kim EH, Oh MH. Clinicopathologic implication of ezrin expression in non-small cell lung cancer. Korean J Pathol. 2012;46(5):470–7. doi: 10.4132/KoreanJPathol.2012.46.5.470. PubMed PMID: 23136574; PubMed Central PMCID: PMCPMC3490123.

97. Aman Wang CL, Zhen Ning2, Wei Gao, Yunpeng Xie, Ningning Zhang, Jinxiao Liang, Faisal S. Abbasi, Qiu Yan, Jiwei Liu. Tumor-associated macrophages promote Ezrin phosphorylation-mediated epithelial-mesenchymal transition in lung adenocarcinoma through FUT4/LeY up-regulation. Oncotarget. 2017;8(17):28247–59.

98. Elliott. VHASAGPAGGPCtaBE. Ezrin regulates focal adhesion and invadopodia dynamics by altering calpain activity to promote breast cancer cell invasion. Molecular Biology of the Cell. 2015;26(19):3464–79.

99. Zhou GZ, Cao FK, Du SW. The apoptotic pathways in the curcumin analog MHMD-induced lung cancer cell death and the essential role of actin polymerization during apoptosis. Biomed Pharmacother. 2015;71:128–34. doi: 10.1016/j.biopha.2015.02.025. PubMed PMID: 25960227.

100. Honggang Zhao YJ, Zuncheng Zhang. Deguelin inhibits the migration and invasion of lung cancer A549 and H460 cells via regulating actin cytoskeleton rearrangement. Int J Clin Exp Pathol. 2015;8(12):15582–90. PubMed PMID: 26884827.

101. Sheng SH, Zhu HL. Proteomic analysis of pleural effusion from lung adenocarcinoma patients by shotgun strategy. Clin Transl Oncol. 2014;16(2):153–7. doi: 10.1007/s12094-013-1054-9. PubMed PMID: 23907289.

102. Menhofer MH, Kubisch R, Schreiner L, Zorn M, Foerster F, Mueller R, et al. The actin targeting compound Chondramide inhibits breast cancer metastasis via reduction of cellular contractility. PLoS One. 2014;9(11):e112542. doi: 10.1371/journal.pone.0112542. PubMed PMID: 25391145; PubMed Central PMCID: PMCPMC4229209.

103. Flamini MI, Fu XD, Sanchez AM, Giretti MS, Garibaldi S, Goglia L, et al. Effects of raloxifene on breast cancer cell migration and invasion through the actin cytoskeleton. J Cell Mol Med. 2009;13(8B):2396–407. doi: 10.1111/j.1582-4934.2008.00505.x. PubMed PMID: 18798864.

104. Jo M, Yun HM, Park KR, Park MH, Lee DH, Cho SH, et al. Anti-cancer effect of thiacremonone through down regulation of peroxiredoxin 6. PLoS One. 2014;9(3):e91508. doi: 10.1371/journal.pone.0091508. PubMed PMID: 24618722; PubMed Central PMCID: PMCPMC3950181.

105. Yun HM, Park KR, Lee HP, Lee DH, Jo M, Shin DH, et al. PRDX6 promotes lung tumor progression via its GPx and iPLA2 activities. Free Radic Biol Med. 2014;69:367–76. doi: 10.1016/j.freeradbiomed.2014.02.001. PubMed PMID: 24512906.

106. Chang XZ, Li DQ, Hou YF, Wu J, Lu JS, Di GH, et al. Identification of the functional role of peroxiredoxin 6 in the progression of breast cancer. Breast Cancer Res. 2007;9(6):R76. doi: 10.1186/bcr1789. PubMed PMID: 17980029; PubMed Central PMCID: PMCPMC2246172.

107. Zhu W, Li GX, Chen HL, Liu XY. The role of eukaryotic translation initiation factor 6 in tumors. Oncol Lett. 2017;14(1):3–9. doi: 10.3892/ol.2017.6161. PubMed PMID: 28693127; PubMed Central PMCID: PMCPMC5494901.

108. Ryo Maeda M, Genichiro Ishii, MD, PhD,Shinya Neri, MD,Kazuhiko Aoyagi, PhD,Hironori Haga, MD, PhD,Hiroki Sasaki, MD, PhD,Kanji Nagai, MD, PhD, and Atsushi Ochiai M, PhD. Circulating CD14+CD204+ Cells Predict Postoperative Recurrence in Non-Small-Cell Lung Cancer Patients. Journal of Thoracic Oncology. 2014;2(9):179–88. PubMed PMID: 24419414.

109. Tian T, Gu X, Zhang B, Liu Y, Yuan C, Shao L, et al. Increased circulating CD14(+)HLA-DR-/low myeloid-derived suppressor cells are associated with poor prognosis in patients with small-cell lung cancer. Cancer Biomark. 2015;15(4):425–32. doi: 10.3233/CBM-150473. PubMed PMID: 25792471.

110. Lobba AR, Forni MF, Carreira AC, Sogayar MC. Differential expression of CD90 and CD14 stem cell markers in malignant breast cancer cell lines. Cytometry A. 2012;81(12):1084–91. doi: 10.1002/cyto.a.22220. PubMed PMID: 23090904.

111. He W, Tong Y, Wang Y, Liu J, Luo G, Wu J, et al. Serum soluble CD14 is a potential prognostic indicator of recurrence of human breast invasive ductal carcinoma with Her2-enriched subtype. PLoS One. 2013;8(9):e75366. doi: 10.1371/journal.pone.0075366. PubMed PMID: 24086515; PubMed Central PMCID: PMCPMC3783397.

112. Grills C, Jithesh PV, Blayney J, Zhang SD, Fennell DA. Gene expression meta-analysis identifies VDAC1 as a predictor of poor outcome in early stage non-small cell lung cancer. PLoS One. 2011;6(1):e14635. doi: 10.1371/journal.pone.0014635. PubMed PMID: 21297950; PubMed Central PMCID: PMCPMC3031508.

113. Azeez JM, Vini R, Remadevi V, Surendran A, Jaleel A, Santhosh Kumar TR, et al. VDAC1 and SERCA3 Mediate Progesterone-Triggered Ca2(+) Signaling in Breast Cancer Cells. J Proteome Res. 2018;17(1):698–709. Epub 2017/12/01. doi: 10.1021/acs.jproteome.7b00754. PubMed PMID: 29185755.

114. Zhang J, Wang J, Xing H, Li Q, Zhao Q, Li J. Down-regulation of FBP1 by ZEB1-mediated repression confers to growth and invasion in lung cancer cells. Molecular and cellular biochemistry. 2016;411(1-2):331–40. Epub 2015/11/08. doi: 10.1007/s11010-015-2595-8. PubMed PMID: 26546081.

115. Li K, Ying M, Feng D, Du J, Chen S, Dan B, et al. Fructose-1,6-bisphosphatase is a novel regulator of Wnt/beta-Catenin pathway in breast cancer. Biomed Pharmacother. 2016;84:1144–9. Epub 2016/10/26. doi: 10.1016/j.biopha.2016.10.050. PubMed PMID: 27780144.

116. Shi L, He C, Li Z, Wang Z, Zhang Q. FBP1 modulates cell metabolism of breast cancer cells by inhibiting the expression of HIF-1alpha. Neoplasma. 2017;64(4):535–42. Epub 2017/05/10. doi: 10.4149/neo 2017 407. PubMed PMID: 28485159.

117. Du S, Guan Z, Hao L, Song Y, Wang L, Gong L, et al. Fructose-bisphosphate aldolase a is a potential metastasis-associated marker of lung squamous cell carcinoma and promotes lung cell tumorigenesis and migration. PLoS One. 2014;9(1):e85804. Epub 2014/01/28. doi: 10.1371/journal.pone.0085804. PubMed PMID: 24465716; PubMed Central PMCID: PMCPmc3900443.

118. Whelan SA, He J, Lu M, Souda P, Saxton RE, Faull KF, et al. Mass spectrometry (LC-MS/MS) identified proteomic biosignatures of breast cancer in proximal fluid. J Proteome Res. 2012;11(10):5034–45. Epub 2012/09/01. doi: 10.1021/pr300606e. PubMed PMID: 22934887; PubMed Central PMCID: PMCPmc3521600.

119. Kawamura M, Endo C, Sakurada A, Hoshi F, Notsuda H, Kondo T. The prognostic significance of eukaryotic elongation factor 1 alpha-2 in non-small cell lung cancer. Anticancer research. 2014;34(2):651–8. Epub 2014/02/11. PubMed PMID: 24510995.

120. Pecorari L, Marin O, Silvestri C, Candini O, Rossi E, Guerzoni C, et al. Elongation Factor 1 alpha interacts with phospho-Akt in breast cancer cells and regulates their proliferation, survival and motility. Molecular cancer. 2009;8:58. Epub 2009/08/04. doi: 10.1186/1476-4598-8-58. PubMed PMID: 19646290; PubMed Central PMCID: PMCPmc2727493.

121. Edmonds BT, Wyckoff J, Yeung YG, Wang Y, Stanley ER, Jones J, et al. Elongation factor-1 alpha is an overexpressed actin binding protein in metastatic rat mammary adenocarcinoma. Journal of cell science. 1996;109 (Pt 11):2705–14. Epub 1996/11/01. PubMed PMID: 8937988.

122. Ihedioha OC, Shiu RP, Uzonna JE, Myal Y. Prolactin-Inducible Protein: From Breast Cancer Biomarker to Immune Modulator-Novel Insights from Knockout Mice. DNA and cell biology. 2016;35(10):537–41. Epub 2016/09/08. doi: 10.1089/dna.2016.3472. PubMed PMID: 27602994.

123. Guetschow ED, Black W, Walsh CM, Furchak JR. Detection of prolactin inducible protein mRNA, a biomarker for breast cancer metastasis, using a molecular beacon-based assay. Analytical and bioanalytical chemistry. 2012;404(2):399–406. Epub 2012/06/14. doi: 10.1007/s00216-012-6162-9. PubMed PMID: 22692591.

124. Clark JW, Snell L, Shiu RP, Orr FW, Maitre N, Vary CP, et al. The potential role for prolactin-inducible protein (PIP) as a marker of human breast cancer micrometastasis. Br J Cancer. 1999;81(6):1002–8. Epub 1999/11/27. doi: 10.1038/sj.bjc.6690799. PubMed PMID: 10576657; PubMed Central PMCID: PMCPmc2362951.

125. He P, Varticovski L, Bowman ED, Fukuoka J, Welsh JA, Miura K, et al. Identification of carboxypeptidase E and gamma-glutamyl hydrolase as biomarkers for pulmonary neuroendocrine tumors by cDNA microarray. Human pathology. 2004;35(10):1196–209. Epub 2004/10/20. PubMed PMID: 15492986.

126. Shubbar E, Helou K, Kovacs A, Nemes S, Hajizadeh S, Enerback C, et al. High levels of gamma-glutamyl hydrolase (GGH) are associated with poor prognosis and unfavorable clinical outcomes in invasive breast cancer. BMC Cancer. 2013;13:47. Epub 2013/02/05. doi: 10.1186/1471-2407-13-47. PubMed PMID: 23374458; PubMed Central PMCID: PMCPmc3576262.

127. Yu CJ, Wang CL, Wang CI, Chen CD, Dan YM, Wu CC, et al. Comprehensive proteome analysis of malignant pleural effusion for lung cancer biomarker discovery by using multidimensional protein identification technology. J Proteome Res. 2011;10(10):4671–82. Epub 2011/08/03. doi: 10.1021/pr2004743. PubMed PMID: 21806062.

128. Dowling P, Clarke C, Hennessy K, Torralbo-Lopez B, Ballot J, Crown J, et al. Analysis of acute-phase proteins, AHSG, C3, CLI, HP and SAA, reveals distinctive expression patterns associated with breast, colorectal and lung cancer. Int J Cancer. 2012;131(4):911–23. Epub 2011/09/29. doi: 10.1002/ijc.26462. PubMed PMID: 21953030.

129. Yi JK, Chang JW, Han W, Lee JW, Ko E, Kim DH, et al. Autoantibody to tumor antigen, alpha 2-HS glycoprotein: a novel biomarker of breast cancer screening and diagnosis. Cancer epidemiology, biomarkers & prevention: a publication of the American Association for Cancer Research, cosponsored by the American Society of Preventive Oncology. 2009;18(5):1357–64. Epub 2009/05/09. doi: 10.1158/1055-9965.epi-08-0696. PubMed PMID: 19423516.

130. Fernandez-Grijalva AL, Aguilar-Lemarroy A, Jave-Suarez LF, Gutierrez-Ortega A, Godinez-Melgoza PA, Herrera-Rodriguez SE, et al. Alpha 2HS-glycoprotein, a tumor-associated antigen (TAA) detected in Mexican patients with early-stage breast cancer. J Proteomics. 2015;112:301–12. Epub 2014/08/12. doi: 10.1016/jjprot.2014.07.025. PubMed PMID: 25106788.

131. Schaub NP, Jones KJ, Nyalwidhe JO, Cazares LH, Karbassi ID, Semmes OJ, et al. Serum proteomic biomarker discovery reflective of stage and obesity in breast cancer patients. Journal of the American College of Surgeons. 2009;208(5):970–8; discussion 8-80. Epub 2009/05/30. doi: 10.1016/jjamcollsurg.2008.12.024. PubMed PMID: 19476873.

132. Cai J, Gu B, Cao F, Liu S. A transferrin-target magnetic/fluorescent dual-mode probe significantly enhances the diagnosis of non-small cell lung cancer. Oncotarget. 2016;7(26):40047–59. Epub 2016/10/26. doi: 10.18632/oncotarget.9482. PubMed PMID: 27223075; PubMed Central PMCID: PMCPmc5129991.

133. Zhang B, Zhang Y, Yu D. Lung cancer gene therapy: Transferrin and hyaluronic acid dual ligand-decorated novel lipid carriers for targeted gene delivery. Oncol Rep. 2017;37(2):937–44. Epub 2016/12/14. doi: 10.3892/or.2016.5298. PubMed PMID: 27959442.

134. Dowling P, Palmerini V, Henry M, Meleady P, Lynch V, Ballot J, et al. Transferrin-bound proteins as potential biomarkers for advanced breast cancer patients. BBA Clin. 2014;2:24–30. Epub 2015/12/18. doi: 10.1016/j.bbacli.2014.08.004. PubMed PMID: 26673961; PubMed Central PMCID: PMCPmc4633920.

135. Xu Q, Zhu M, Yang T, Xu F, Liu Y, Chen Y. Quantitative assessment of human serum transferrin receptor in breast cancer patients pre- and post-chemotherapy using peptide immunoaffinity enrichment coupled with targeted proteomics. Clin Chim Acta. 2015;448:118–23. Epub 2015/06/23. doi: 10.1016/j.cca.2015.05.022. PubMed PMID: 26096257.

136. Okudela K, Katayama A, Woo T, Mitsui H, Suzuki T, Tateishi Y, et al. Proteome analysis for downstream targets of oncogenic KRAS–the potential participation of CLIC4 in carcinogenesis in the lung. PLoS One. 2014;9(2):e87193. Epub 2014/02/08. doi: 10.1371/journal.pone.0087193. PubMed PMID: 24503901; PubMed Central PMCID: PMCPmc3913595.

137. Chiang PC, Chou RH, Chien HF, Tsai T, Chen CT. Chloride intracellular channel 4 involves in the reduced invasiveness of cancer cells treated by photodynamic therapy. Lasers in surgery and medicine. 2013;45(1):38–47. Epub 2013/01/17. doi: 10.1002/lsm.22112. PubMed PMID: 23322262.

138. Yang H, Chen J, Yang J, Qiao S, Zhao S, Yu L. Cyclophilin A is upregulated in small cell lung cancer and activates ERK1/2 signal. Biochemical and biophysical research communications. 2007;361(3):763–7. Epub 2007/08/07. doi: 10.1016/j.bbrc.2007.07.085. PubMed PMID: 17678621.

139. Campa MJ, Wang MZ, Howard B, Fitzgerald MC, Patz EF, Jr. Protein expression profiling identifies macrophage migration inhibitory factor and cyclophilin a as potential molecular targets in non-small cell lung cancer. Cancer Res. 2003;63(7):1652–6. Epub 2003/04/03. PubMed PMID: 12670919.

140. Tamesa MS, Kuramitsu Y, Fujimoto M, Maeda N, Nagashima Y, Tanaka T, et al. Detection of autoantibodies against cyclophilin A and triosephosphate isomerase in sera from breast cancer patients by proteomic analysis. Electrophoresis. 2009;30(12):2168–81. Epub 2009/07/08. doi: 10.1002/elps.200800675. PubMed PMID: 19582718.

141. Chevalier F, Depagne J, Hem S, Chevillard S, Bensimon J, Bertrand P, et al. Accumulation of cyclophilin A isoforms in conditioned medium of irradiated breast cancer cells. Proteomics. 2012;12(11):1756–66. Epub 2012/05/25. doi: 10.1002/pmic.201100319. PubMed PMID: 22623065.

142. He P, Naka T, Serada S, Fujimoto M, Tanaka T, Hashimoto S, et al. Proteomics-based identification of alpha-enolase as a tumor antigen in non-small lung cancer. Cancer science. 2007;98(8):1234–40. Epub 2007/05/18. doi: 10.1111/j.1349-7006.2007.00509.x. PubMed PMID: 17506794.

143. Chang YS, Wu W, Walsh G, Hong WK, Mao L. Enolase-alpha is frequently down-regulated in non-small cell lung cancer and predicts aggressive biological behavior. Clin Cancer Res. 2003;9(10 Pt 1):3641–4. Epub 2003/09/25. PubMed PMID: 14506152.

144. Tu SH, Chang CC, Chen CS, Tam KW, Wang YJ, Lee CH, et al. Increased expression of enolase alpha in human breast cancer confers tamoxifen resistance in human breast cancer cells. Breast Cancer Res Treat. 2010;121(3):539–53. Epub 2009/08/06. doi: 10.1007/s10549-009-0492-0. PubMed PMID: 19655245.

145. Zhao H, Jiao Y, Zhang Z. Deguelin inhibits the migration and invasion of lung cancer A549 and H460 cells via regulating actin cytoskeleton rearrangement. Int J Clin Exp Pathol. 2015;8(12):15582–90. Epub 2016/02/18. PubMed PMID: 26884827; PubMed Central PMCID: PMCPmc4730040.

146. Zhang D, Zhang JY, Dai SD, Liu SL, Liu Y, Tang N, et al. Co-expression of delta-catenin and RhoA is significantly associated with a malignant lung cancer phenotype. Int J Clin Exp Pathol. 2014;7(7):3724–32. Epub 2014/08/15. PubMed PMID: 25120748; PubMed Central PMCID: PMCPmc4128983.

147. Yang X, Zheng F, Zhang S, Lu J. Loss of RhoA expression prevents proliferation and metastasis of SPCA1 lung cancer cells in vitro. Biomed Pharmacother. 2015;69:361–6. Epub 2015/02/11. doi: 10.1016/j.biopha.2014.12.004. PubMed PMID: 25661383.

148. Lei R, Tang J, Zhuang X, Deng R, Li G, Yu J, et al. Suppression of MIM by microRNA-182 activates RhoA and promotes breast cancer metastasis. Oncogene. 2014;33(10):1287–96. Epub 2013/03/12. doi: 10.1038/onc.2013.65. PubMed PMID: 23474751.

149. Lee HK, Choung HW, Yang YI, Yoon HJ, Park IA, Park JC. ODAM inhibits RhoA-dependent invasion in breast cancer. Cell biochemistry and function. 2015;33(7):451–61. Epub 2015/09/12. doi: 10.1002/cbf.3132. PubMed PMID: 26358398.

150. Jang SM, An JH, Kim CH, Kim JW, Choi KH. Transcription factor FOXA2-centered transcriptional regulation network in non-small cell lung cancer. Biochemical and biophysical research communications. 2015;463(4):961–7. Epub 2015/06/21. doi: 10.1016/j.bbrc.2015.06.042. PubMed PMID: 26093302.

151. Prasad S, Soldatenkov VA, Srinivasarao G, Dritschilo A. Identification of keratins 18, 19 and heat-shock protein 90 beta as candidate substrates of proteolysis during ionizing radiation-induced apoptosis of estrogen-receptor negative breast tumor cells. Int J Oncol. 1998;13(4):757–64. Epub 1998/09/15. PubMed PMID: 9735406.

152. Song Q, Xu Y, Yang C, Chen Z, Jia C, Chen J, et al. miR-483-5p promotes invasion and metastasis of lung adenocarcinoma by targeting RhoGDI1 and ALCAM. Cancer Res. 2014;74(11):3031–42. Epub 2014/04/09. doi: 10.1158/0008-5472.can-13-2193. PubMed PMID: 24710410.

153. Ronneburg H, Span PN, Kantelhardt E, Dittmer A, Schunke D, Holzhausen HJ, et al. Rho GDP dissociation inhibitor alpha expression correlates with the outcome of CMF treatment in invasive ductal breast cancer. Int J Oncol. 2010;36(2):379–86. Epub 2010/01/01. PubMed PMID: 20043072.

154. Jiang H, Wu L, Mishra M, Chawsheen HA, Wei Q. Expression of peroxiredoxin 1 and 4 promotes human lung cancer malignancy. American journal of cancer research. 2014;4(5):445–60. Epub 2014/09/19. PubMed PMID: 25232487; PubMed Central PMCID: PMCPmc4163610.

155. Kim JH, Bogner PN, Baek SH, Ramnath N, Liang P, Kim HR, et al. Up-regulation of peroxiredoxin 1 in lung cancer and its implication as a prognostic and therapeutic target. Clin Cancer Res. 2008;14(8):2326–33. Epub 2008/04/17. doi: 10.1158/1078-0432.ccr-07-4457. PubMed PMID: 18413821.

156. O'Leary PC, Terrile M, Bajor M, Gaj P, Hennessy BT, Mills GB, et al. Peroxiredoxin-1 protects estrogen receptor alpha from oxidative stress-induced suppression and is a protein biomarker of favorable prognosis in breast cancer. Breast Cancer Res. 2014;16(4):R79. Epub 2014/07/12. doi: 10.1186/bcr3691. PubMed PMID: 25011585; PubMed Central PMCID: PMCPmc4226972.

157. Fujimoto J, Kadara H, Garcia MM, Kabbout M, Behrens C, Liu DD, et al. G-protein coupled receptor family C, group 5, member A (GPRC5A) expression is decreased in the adjacent field and normal bronchial epithelia of patients with chronic obstructive pulmonary disease and non-small-cell lung cancer. Journal of thoracic oncology: official publication of the International Association for the Study of Lung Cancer. 2012;7(12):1747–54. Epub 2012/11/17. doi: 10.1097/JTO.0b013e31826bb1ff. PubMed PMID: 23154545; PubMed Central PMCID: PMCPmc3622592.

158. Chatterji B, Borlak J. A 2-DE MALDI-TOF study to identify disease regulated serum proteins in lung cancer of c-myc transgenic mice. Proteomics. 2009;9(4):1044–56. Epub 2009/01/31. doi: 10.1002/pmic.200701135. PubMed PMID: 19180532.

159. Jiao C, Cui L, Ma A, Li N, Si H. Elevated Serum Levels of Retinol-Binding Protein 4 Are Associated with Breast Cancer Risk: A Case-Control Study. PLoS One. 2016;11(12):e0167498. Epub 2016/12/22. doi: 10.1371/journal.pone.0167498. PubMed PMID: 28002423; PubMed Central PMCID: PMCPmc5176270.

160. Pietrowska M, Jelonek K, Michalak M, Ros M, Rodziewicz P, Chmielewska K, et al. Identification of serum proteome components associated with progression of non-small cell lung cancer. Acta biochimica Polonica. 2014;61(2):325–31. Epub 2014/05/30. PubMed PMID: 24872961.

161. Chung L, Moore K, Phillips L, Boyle FM, Marsh DJ, Baxter RC. Novel serum protein biomarker panel revealed by mass spectrometry and its prognostic value in breast cancer. Breast Cancer Res. 2014;16(3):R63. Epub 2014/06/18. doi: 10.1186/bcr3676. PubMed PMID: 24935269; PubMed Central PMCID: PMCPmc4095593.

162. Chen WH, Yin HL, Chen CJ. Anti-beta2-glycoprotein I antibody and cerebellar ataxia in breast cancer. Lupus. 2012;21(4):460–2. Epub 2012/03/20. doi: 10.1177/0961203312437436. PubMed PMID: 22427365.

163. Matboli M, Eissa S, Said H. Evaluation of histidine-rich glycoprotein tissue RNA and serum protein as novel markers for breast cancer. Med Oncol. 2014;31(4):897. Epub 2014/02/26. doi: 10.1007/s12032-014-0897-4. PubMed PMID: 24567057.

164. Lysov Z, Swystun LL, Kuruvilla S, Arnold A, Liaw PC. Lung cancer chemotherapy agents increase procoagulant activity via protein disulfide isomerase-dependent tissue factor decryption. Blood coagulation & fibrinolysis: an international journal in haemostasis and thrombosis. 2015;26(1):36–45. Epub 2014/06/10. doi: 10.1097/mbc.0000000000000145. PubMed PMID: 24911456.

165. Wise R, Duhachek-Muggy S, Qi Y, Zolkiewski M, Zolkiewska A. Protein disulfide isomerases in the endoplasmic reticulum promote anchorage-independent growth of breast cancer cells. Breast Cancer Res Treat. 2016;157(2):241–52. Epub 2016/05/11. doi: 10.1007/s10549-016-3820-1. PubMed PMID: 27161215; PubMed Central PMCID: PMCPmc5662471.

